# Molecular dissection of protein complexes isolated from sections of human brain

**DOI:** 10.64898/2026.04.11.717886

**Authors:** Tarick J. El-Baba, Corinne A. Lutomski, Jack L. Bennett, Sophie A. S. Lawrence, Sean A. Burnap, Frances I. Butroid, Olivia B. Ramsay, Titas Radzevičius, Di Wu, Haigang Song, Kenny L. Chan, Lyonna F. Parise, Eric M. Parise, Weston B. Struwe, James W. Murrough, Scott J. Russo, Carol V. Robinson

## Abstract

Molecular studies of brain receptors and transporters have typically relied on recombinant systems, limiting insight into their organization in native tissue. Here, we develop nanobody-based immunoprecipitation coupled with native mass spectrometry to interrogate endogenous protein assemblies from post-mortem mouse and human brain sections. We exemplify our approach by characterizing the synaptic proteins VGluT1 and mGluR2. From a single mouse brain, we discover mGluR2/3 heterodimers, alongside mGluR2 homodimers. Considering regions of human brain heterodimeric mGluR2/3 is highly abundant in the OFC and sgACC (∼70% and 50%, respectively) and forms regional-specific interactions with additional synaptic proteins. In a modest cohort of biobanked human tissue, associated with depression and suicide, we find increased mGluR2/3 in the OFC. Consistent with this, mice exhibit similar associations between heterodimer levels and stress-susceptibility. Overall, our approach provides a direct means for establishing molecular-behavioural links at the level of receptor organization in brain.

## Introduction

The brain harbours a remarkably diverse repertoire of protein assemblies, whose region-specific distribution and subunit composition govern essential neural functions. A central feature of many psychiatric disorders is dysfunctional glutamate signaling in frontal cortical regions of the brain, which is broadly controlled by ion channels, transporters, and G protein-coupled receptors (GPCRs).^1^ Recent cryoelectron microscopy (cryoEM) studies have exemplified the complexity of ion channels by revealing differential subunit organizations of glutamate-gated AMPA^2–5^ and NMDA receptors,^6,7^ and heteromeric GABA(A)^8,9^ receptors. CryoEM structures of brain-isolated AMPA and GABA(A) receptors have revealed that these endogenous assemblies have distinct subunit compositions which confer differential protein interactions and unique pharmacological profiles. These structures have largely been derived from pooled murine or human tissue. While informative for defining native complex architecture, they are not able to resolve individual-level variation in receptor composition and interaction that is needed to align with behavioural phenotypes and guide initiatives in precision medicine. Accordingly, elucidating the molecular organization of endogenous, membrane-embedded regulators of neuronal activity is of considerable importance.

Considering steps in the glutamatergic signalling system, glutamate is first loaded into synaptic vesicles by vesicular glutamate transporters (VGluTs) and released via exocytosis from the presynaptic terminal. It then activates metabotropic glutamate receptors (mGluRs), class C GPCRs that are highly relevant to synaptic processing.^10,11^ Group II mGluRs (mGluR2 and mGluR3) are widely expressed in the brain and are considered key regulators of excitatory neurotransmission. All functional mGluRs are obligate dimers^10,12^ and have the possibility to form heterodimers that have distinct functional and pharmacological properties.^13–16^ mGluR heterodimers have been studied as isolated proteins,^17–20^ ectopically expressed in model cell lines,^21–23^ in cultured neuronal cells,^24^ in mouse brain slices,^15^ and in membrane fractions.^14,25^ Whether homo- and heterodimeric assemblies co-exist in human brain, how their relative abundance varies across brain regions, and if they are organized within larger protein complexes, remains largely unknown.

Proteomics and transcriptomics provide powerful insights into protein and gene abundance in brain;^26–30^ yet central unresolved questions concern the organization of proteoforms into complexes within a regional context and across varying disease states. Answering these questions demands methods capable of resolving and preserving the composition and interactions of low-abundance endogenous receptor complexes, directly from brain tissue, with molecular precision. Native mass spectrometry (nMS) and native top-down mass spectrometry (nTDMS) have recently enabled intact mass measurements, subunit-level identification, and post-translational modification (PTM) status of endogenous protein complexes, including novel assemblies detected directly from cell membranes.^31,32^ In nMS, a protein complex of interest is introduced directly into the mass spectrometer under non-denaturing conditions; stoichiometry and binding interactions, that lead to distinct mass changes, can be readily detected.^33–37^ Where mass analysis alone is insufficient, nTDMS can be used hierarchically to dissociate protein complexes into constituent subunits, bound ligands, and peptide-level fragments for sequence analysis, all within the mass spectrometer.^31,38–42^ However, these approaches have typically been limited to recombinant proteins or abundant complexes in cells or tissues.^34,35,43,44^ nMS and nTDMS of endogenous synaptic receptors, isolated directly from brain tissue, has remained beyond reach.

Here, we address this gap by presenting a nanobody (Nb)-based immunoprecipitation platform enabling capture of multiple native synaptic receptor complexes from single brain tissue specimens in less than 24 h. We demonstrate that our strategy recovers transporters and receptors from minute quantities of anatomically dissected mouse and post-mortem human tissue at levels suitable for comprehensive MS-based characterization. By integrating nMS, nTDMS, glycoproteomics, bound-lipid analysis, and chemical crosslinking, we define the composition, stoichiometry, glycosylation and palmitoylation state, and protein interactions of VGluT1- and mGluR2-containing complexes with unprecedented resolution. We unambiguously identify mGluR2/3 heterodimers in both mouse and human brain. Moreover, we demonstrate distinct species- and region-specific heterodimer abundances, and define region-specific protein interactions, including a direct association between CRMP2 and the mGluR3 subunit of mGluR2/3 heterodimers in the sgACC. Applying this framework to biobanked OFC tissue from individuals with depression that died by suicide, we find that heterodimer levels are associated with end-stage depression.

## Results

### Isolation of native glutamate transporters and receptors from individual brains

To probe the organization of protein complexes in brain, we devised a Nb^45–47^-based immunoprecipitation strategy capable of capturing multiple synaptic assemblies from single brain sections for interrogation by nMS and nTDMS (**Figure 1A**). Critically, the homogenous molecular composition and small size of Nbs eliminates the need to disrupt Nb-target interactions prior to nMS and nTDMS, thereby preserving labile assembly states and interactions through downstream analysis. For retrieval of VGluT1 and mGluR2 from single sections described here, the entire process can be completed in under 24 h following tissue harvest.

**Figure 1.**
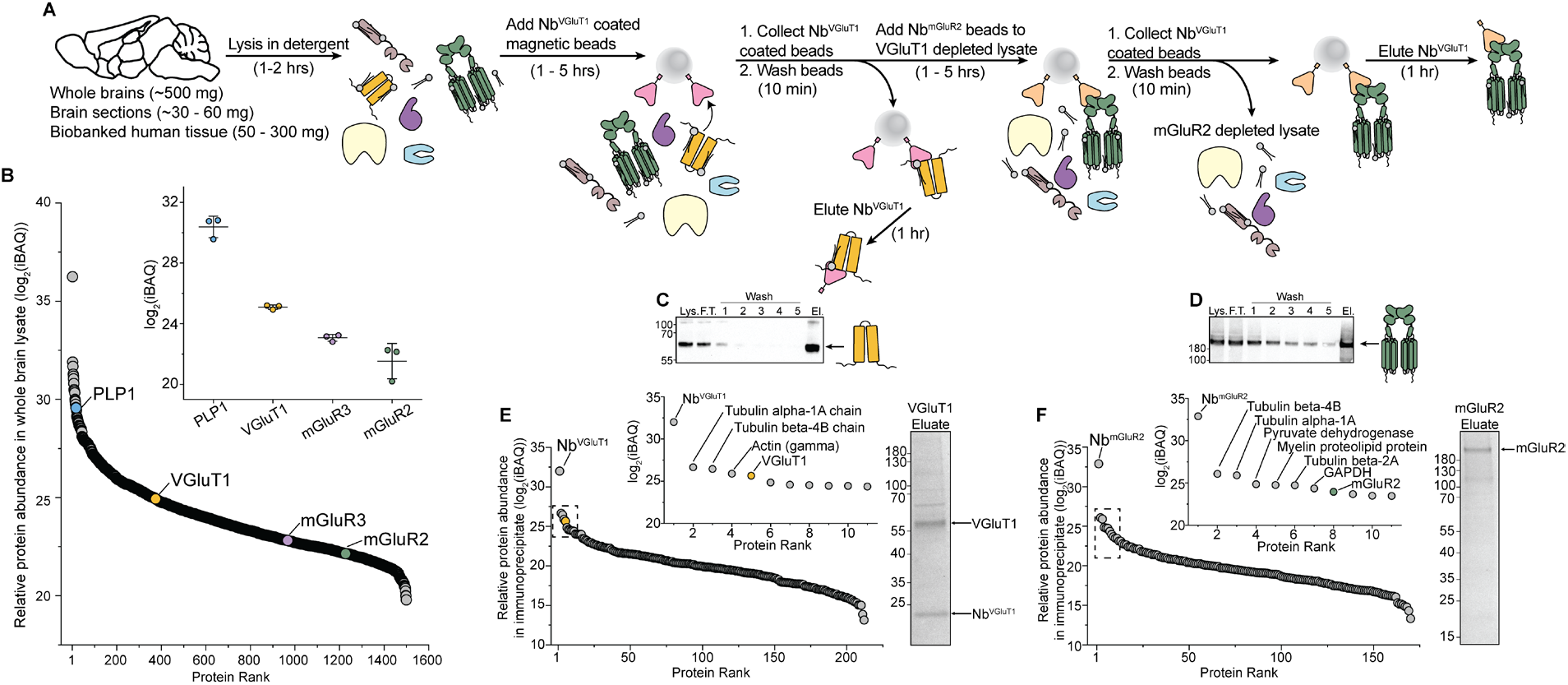
A Nb-based immunoprecipitation strategy for recovering multiple synaptic protein complexes from individual brains. (A) The overall workflow depicting immunoprecipitation of VGluT1 and mGluR2. Briefly, a mouse brain (∼400 mg) is lysed in detergent-containing buffer. The lysate is then incubated with Nb^VGluT1^-coated anti-FLAG magnetic agarose beads. Beads containing captured VGluT1 are collected using a magnet, washed in detergent-containing buffer to release non-specifically associated proteins and lipids; the VGluT1-depleted lysate is collected. The Nb^VGluT1^-coated beads, bound VGluT1, and associated interactors (lipids and proteins) are eluted using FLAG peptide. Next, the VGluT1-depleted lysate is incubated with beads containing Nb^mGluR2^ and the process is repeated to immunocapture mGluR2. The entire procedure can be completed rapidly (less than 24 h herein) and is dependent on the k_on_/k_off_ parameters of the Nb. Numbers in parentheses indicates approximate time for each step. (B) Ranked protein abundance plot depicting the distribution of proteins and their relative abundances in a single C57BL6 mouse brain lysate. PLP1: Proteolipid Protein 1 (myelin), VGluT1, mGluR2 and mGluR3 are highlighted (light blue, orange, purple, and green respectively). Inset shows the relative abundances of each protein in n=3 independent mouse brain lysates. (C) Immunoblot tracking the immunoprecipitation of VGluT1 using Nb^VGluT1^. (D) Immunoblot following the immunoprecipitation of mGluR2 using Nb^mGluR2^. Lys., Lysate; F.T., Flow-through; El., Elution. (E) Plot of relative protein abundance (log_2_(iBAQ)) relative to protein rank to identify and quantify the proteins that co-immunoprecipitated with VGluT1. (F) Plot of proteomic abundance relative to protein rank identifying and quantifying the proteins which coimmunoprecipitate with mGluR2.

We first optimized our approach using whole C57BL6 mouse brains. Shotgun proteomics of whole brain lysates identified approximately ∼1,600 proteins, and confirmed that VGluT1 and mGluR2 are present but at low abundance relative to cytoskeletal proteins and metabolic enzymes (**Figure 1B**). Using an optimized approach, we first immunoprecipitated VGluT1, then from the same lysate mGluR2 using FLAG-tagged Nbs (Nb^VGluT1^;Ref. ^48^ Ref. and Nb^mGluR2^; Ref.^49^ and WO2016001417A1). Western blotting confirmed enrichment of both VGluT1 and mGluR2 (**Figure 1C,D**). Proteomic analysis of the enriched immunoprecipitates showed that both targets were among the ten most abundant proteins, below only the Nbs and common structural proteins (**Figure 1E,F**).

Turning to nMS, a ∼1 µM eluate of immunoprecipitated murine VGluT1, stabilized in detergent micelles, was introduced into a modified Orbitrap Ascend mass spectrometer^31^. Using optimised conditions^31,39^ we successfully liberated the transporter from micelles and obtained a well-resolved charge state distribution corresponding to an average molecular weight of 82,282 ± 45 Da (**Figure 2A**). We applied nTDMS^31,50,51^, and generated a spectrum containing abundant fragment ions assigned to murine VGluT1 and Nb^VGluT1^ (**Figure 2B**). This mass is assigned to a noncovalent assembly between the Nb^VGluT1^ (17,782 Da) and murine VGluT1 (sequence mass 61,637 Da), with residual unexplained mass of 2,863 Da. Glycoproteomic analysis identified a single occupied glycosylation site at Asn-93, which harbored a sialylated tri-antennary N-glycan (GlcNAc_5_Hex_5_Fuc_1_NeuAc_1_, MW = 2,262 Da) (**Figure 2A inset**). To account for the remaining unexplained mass, we applied nTDMS under conditions suitable for releasing non-covalently bound ligands without breaking the peptide backbone^52^ (**Figure 2C**). Two abundant peaks, consistent with lysophosphatidylinositol (22:4) and phosphatidylinositol (32:0) were observed amid other low-abundance features. The residual mass can be attributed to multiple lipids bound to murine VGluT1. Other high molecular weight peaks, unresolvable in nMS spectra, can be accounted for by multiple PI, LPI, and other lipid bound states. Phospholipid-like densities have been found in cryoEM structures of VGluTs;^53,54^ it is plausible that the released lipids play a structural role in the glutamate loading into synaptic vesicles.

**Figure 2.**
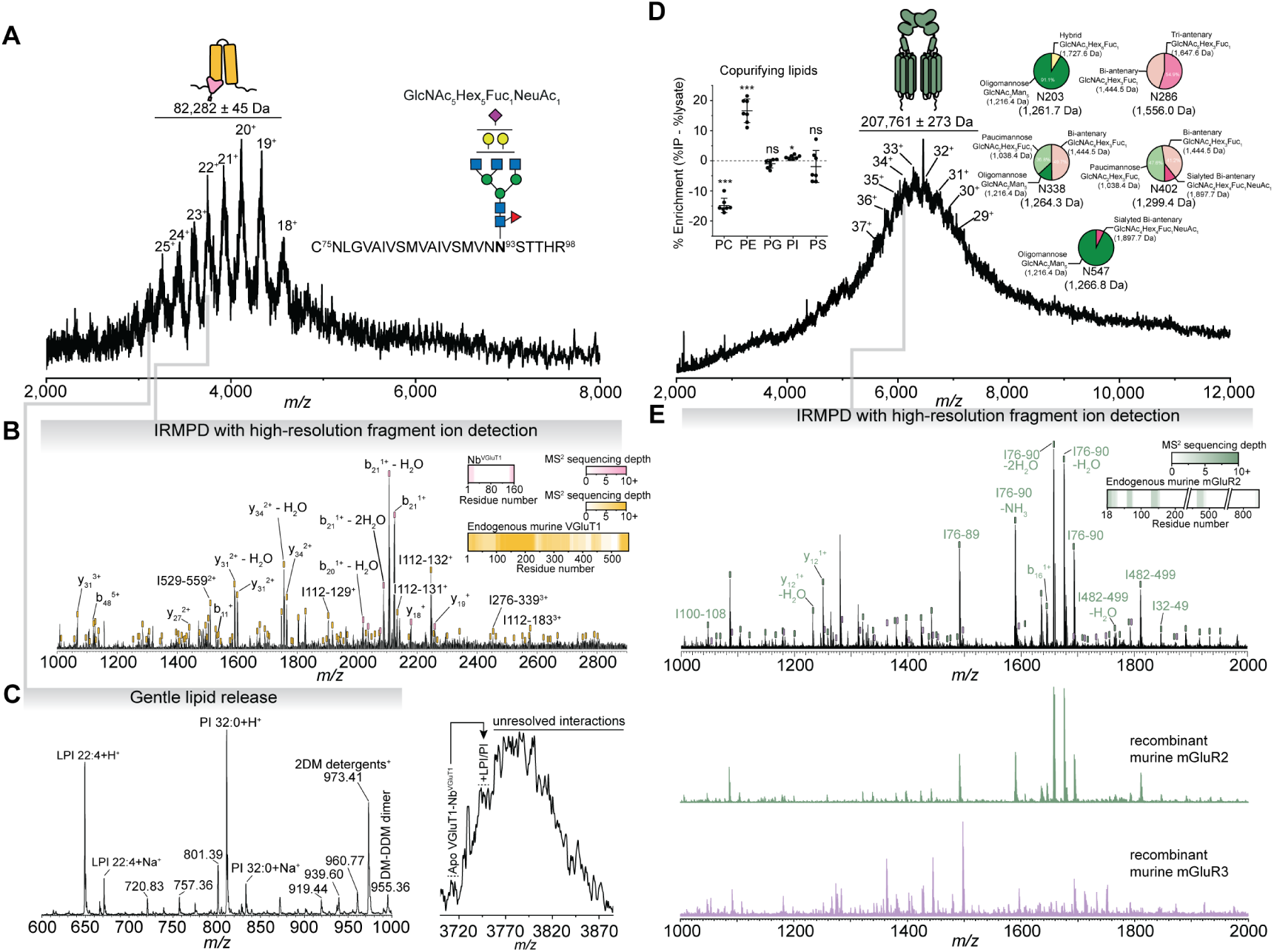
Native mass spectrometry reveals the organization, sequence, and interactions of multiple endogenous synaptic complexes from a single mouse brain. (A) Native mass spectrum of VGluT1 immunoprecipitated from a single mouse brain. Inset shows the position of the only identified N-glycan by glycoproteomics (a paucimannose on Asn93 in the extracellular loop). (B) nTDMS spectrum and inset sequence depth of murine VGluT1 (and Nb^VGluT1^), higher sequencing depth denoted by color intensity. (C) nTDMS spectrum (linear ion trap detection) collected at low activation energies to promote the release of non-covalently bound lipids. Two peaks matched the predicted molecular weight (within 0.2 Da) of lysophosphatidylinositol (LPI) (22:4) and phosphatidylinositol (PI) (32:0). An overlay of the expected peak positions on the native mass spectrum (exemplified with 22^+^ peak) for lipid complexes of LPI or PI with VGluT1-Nb^VGluT1^. (D) Native mass spectrum of mGluR2 from a single mouse brain. Site-specific N-glycoforms (pie charts) with weighted average molecular weights indicated in brackets. Differences in lipid class abundance (strip plot percentages) between mGluR2 immunoprecipitates and total brain lysates for five major phospholipid classes. Values represent IP% − lysate% for each biological replicate (n=7), horizontal bars indicate group means, and error bars the standard deviation. Statistical comparisons were performed using one-sample paired t-tests against a null hypothesis of zero difference, with Holm–Bonferroni correction for five comparisons. *p<0.05, **p<0.01, ***p<0.001. PC, phosphatidylcholine; PE, phosphatidylethanolamine; PG, phosphatidylglycerol; PI, phosphatidylinositol; PS, phosphatidylserine. (E) nTDMS spectrum of murine mGluR2 (top, black) with inset sequencing depth. Higher sequencing depth denoted with more intense coloring. nTDMS spectra of recombinant mouse mGluR2 and mGluR3 dimers (green and purple, respectively) used to build libraries of fragment ions for spectral matching.

We next applied this workflow to the murine mGluR2 isolated from the same brain lysate after VGluT1 depletion. We recorded a native mass spectrum, observing a charge state distribution between *m/z* 5800 – 7500 corresponding to complexes with molecular weights 207,761 ± 273 Da (**Figure 2D** and **Figure S1)**, consistent with dimeric mGluR2. This complex is approximately 19 kDa heavier than the anticipated sequence mass of dimeric mGluR2 (197,863 Da), suggesting extensive N-glycosylation; this assumption is consistent with broad peaks that are characteristic of heterogenous glycosylation in nMS.^55^ Glycoproteomic analysis confirmed complete N-glycan occupancy at all five sequons (**Figure 2D right inset**); site-specific analysis further showed that each sequon carried no more than three distinct glycoforms, spanning paucimannose (GlcNAc_2_Hex_2_Fuc_1_), oligomannose (Man_5_GlcNAc_2_), and bi- or tri-antennary complex-type glycans (GlcNAc_3-5_Hex_3_Fuc_1_). Using the site-specific weighted average glycan masses, N-glycans account for ∼13,294 Da of the residual mass. To account for the remaining ∼6.5 kDa, we next explored the possibility that mGluR2 forms extensive association with lipids from the membrane. Attempts to release lipids bound to mGluR2 via nTDMS yielded no signatures of cationic phospholipids; we instead turned to our native MS-guided lipidomic workflow (**Figure 2D left inset**).^56^ The abundances of neutral PE and PI lipids were significantly enriched with mGluR2 relative to lysates, whereas cationic PC lipids were significantly downregulated. While not precluding them from interacting with mGluR2, anionic PS and neutral PG lipid abundances were not significantly different than their abundance in brain lysates. We assign the remaining mass to neutral or anionic phospholipids from the membrane retained noncovalently on mGluR2.

To generate sequence information and confirm our assignment, we subjected a subset of our tentatively assigned mGluR2 dimer peaks to nTDMS (**Figure 2E, top**). Initially, only ∼10 of the abundant peaks in the nTDMS spectrum could be assigned to fragments of mGluR2 by database searching.^38^ To assign the remaining peaks in the spectrum, we expressed and purified recombinant murine mGluR2 and subjected it to nTDMS to construct a reference fragment ion library (**Figure 2E, middle**). Using this library, we assigned a substantial fraction of the peaks in **Figure 2E (top)** to murine mGluR2. The majority of the fragment ions originate from peptide bond cleavage in the extracellular Venus Fly Trap domain^49^ (residues 18 to ∼500) and the C-terminus (**Figure 2E, inset**). ∼50 low abundance peaks could not be matched to mGluR2, but instead matched to library spectra generated from recombinant murine mGluR3 dimers (**Figure 2E, bottom**). This was unexpected but could potentially be explained by cross reactivity of the Nb^mGluR2^. To explore this possibility, we used nMS to examine Nb^mGluR2^ binding to all combinations of recombinant dimers: mGluR2/2, mGluR3/3 and mGluR2/3 (**Figure S2-4**). Nb^mGluR2^ binding was only observed when mGluR2 was present, consistent with the high specificity of this nanobody (WO2016001417A1).^49^ Our nTDMS data therefore strongly suggest that our immunoprecipitation enriched for both mGluR2/2 homodimers and mGluR2/3 heterodimers, the latter appear to be prevalent in the murine brain. This prompted us to develop a quantification strategy to explore the regional distribution of endogenous heterodimers across the brain.

### Regional distribution of mGluR2/3 heterodimers across the mouse brain

To map the prevalence of mGluR2/3 heterodimers across the murine brain, cerebral cortex, hippocampus, and cerebellum were dissected from a single C57BL6 adult mouse brain and subjected to parallel Nb^mGluR2^ immunoprecipitations. Each region underpins distinct functions: cognition/perception, memory, and voluntary movement, respectively, and together represent a broad sampling of neuroanatomically and functionally diverse tissue. We analyzed ∼100 - 500 nM preparations of cortical, hippocampal, and cerebellar mGluR2 by nMS (**Figure 3A**). In all cases, we could discern dimer peaks near *m/z* 6000. In hippocampal preparations, the small size of this tissue (<30 mg) yielded spectra with slightly reduced signal-to-noise, but still populated by peaks corresponding to dimeric mGluR. Considering nTDMS, in all cases there was evidence for peaks corresponding to mGluR3 amid the predominant mGluR2 fragment ions (**Figure 3A insets**). Together, these results indicate that mGluR2/3 heterodimers are present across neuroanatomically diverse regions of murine brain.

**Figure 3.**
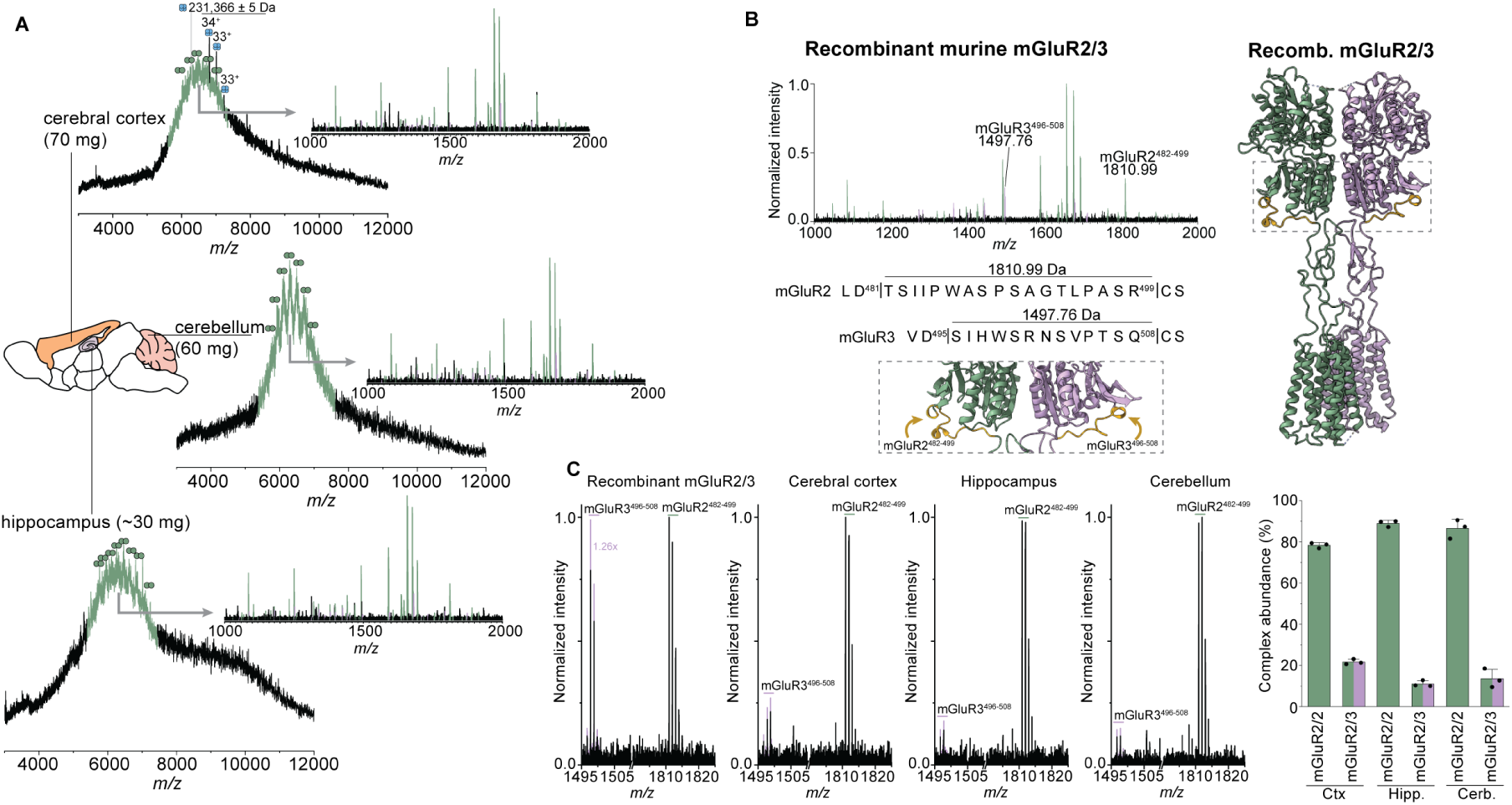
Distribution of endogenous mGluR2/3 heterodimers in neuroanatomically-defined regions in the mouse brain. (A) native mass spectra of mGluR2 immunoprecipitates from cerebral cortex, hippocampus, and cerebellum from n=3 adult C57BL6 mice. Inset show nTDMS spectra, with arrows depicting the region selected for fragmentation and annotations indicating mGluR2 and mGluR3-specific fragments. (B) nTDMS spectrum of recombinant murine mGluR2/3, with mGluR2^482–499^ and mGluR3^495–508^ labelled. The locations of these fragments are shown in gold in the inset and right cryoEM structure (PDB 8JD2).^57^ (C) nTDMS spectra of recombinant murine mGluR2/3 illustrating that a 1:1 heteromeric complex produced mGluR2^76–91^:mGluR3^83–98^ at an intensity ratio 1.26:1 (calibration baseline). These peaks are present in nTDMS spectra from cortical, hippocampal, and cerebellar mGluR2 preparations, albeit at differing intensities. Bar plot depicts the relative abundances of mGluR2/2 and mGluR2/3 across the dissected regions from n=3 C57BL6 adult mice. Bar heights denote the mean and error bars the standard error of the mean.

We reasoned that the relative intensities of specific fragment ions, in the same nTDMS spectrum, could be used to determine the relative proportions of endogenous mGluR2/2 homodimers and mGluR2/3 heterodimers present within a given dimer peak (i.e., complex-specific quantification). To identify suitable reporter ions, we searched nTDMS spectra of recombinant mGluR2 dimers and recombinant mGluR3 dimers for fragment peaks consistently present in endogenous nTDMS spectra. Using a combination of computational analysis^38^ and library matching, we identified two peaks that originate from analogous locations in the lower lobe of the Venus Fly Trap domain of mGluR2 and mGluR3 (**Figure 3B**). In recombinant mGluR2/3 preparations, where the subunits are stoichiometrically equivalent, these two peaks (mGluR2^482–499^ and mGluR3^495–508^) appear in a signal intensity ratio of 1.26:1 (mGluR2:mGluR3), establishing a calibration baseline for a 50:50 mixture of mGluR2/2 and mGluR2/3 (**Figure 3C**). These peaks were found in all nTDMS spectra generated by fragmenting individual dimer peaks from endogenous mGluR2 immunoprecipitation, albeit at different intensity levels depending on brain region (**Figure 3C**). After accounting for calibrated responses of mGluR2^482–499^ and mGluR3^495–508^, the cerebral cortex had the highest heterodimer content (22%); the homodimer:heterodimer ratios in hippocampus (89:11) and cerebellum (86:14) were similar, but lower than in cerebral cortex.

### Human cortical regions have high levels of mGluR2/3 heterodimers

We next sought to understand if heterodimers are also distributed within functionally and neuroanatomically distinct regions in human prefrontal cortex. We focused the orbitofrontal cortex (OFC/Brodmann area 11) and the subgenual anterior cingulate cortex (sgACC/Broadmann area 25); prefrontal regions that serve different but important roles cognitive processes.^1,58^ Engagement of the OFC is strongly associated with reward valuation, decision making and emotion,^1,59,60^ and is found to be hypoactive in severe depression.^61^ While the sgACC also plays a major role in mood regulation, hyperactivity is associated with psychiatric illness.^1,62,63^ Notably, the sgACC is the target of deep brain stimulation for treatment-resistant depression.^64,65^

To probe regional difference of mGluR2/3 in human brain we obtained ∼150 mg of biobanked fresh frozen tissue from sgACC (n=4) (**Table S1**) and OFC (n=7) (**Table S2**) and immunoprecipitated mGluR2. nMS analysis of ∼1 µM preparations from the OFC revealed two charge state distributions (**Figure 4A**): an abundant distribution centered near *m/z* 6800 which corresponds to dimeric mGluR2 and, a sharp series of peaks near *m/z* 9000 corresponding to a complex of mass 421,418 ± 131 Da. SDS-PAGE analyses of an OFC preparation revealed a distinct ∼200 kDa band for mGluR2 and an intense band near 40 kDa which was excised and subjected to proteomic analysis, identifying glutamine synthetase (GS) (**Figure 4A inset and Table S3**). GS is a decameric enzyme that is responsible for the cytosolic conversion of glutamate to glutamine.^67^ The ∼421 kDa complex is consistent with GS_10-mer_. GS has also co-immunoprecipitated with endogenous NMDA receptors;^6^ together with the observation here, suggests that glutamate sensing and its metabolism are coupled. Although the masses and relative abundance of peptide fragments differed from murine to human, both mGluR2 and mGluR3 fragments were present in nTDMS spectra, indicating heterodimeric mGluR2/3 is present in OFC preparations. Interestingly, we also observed fragment ions corresponding to palmitoylation at Cys817 or Cys866 on the mGluR3 C-terminal tail, indicating that endogenous mGluR2/3 heterodimers are palmitoylated (**Figure S5**).

**Figure 4.**
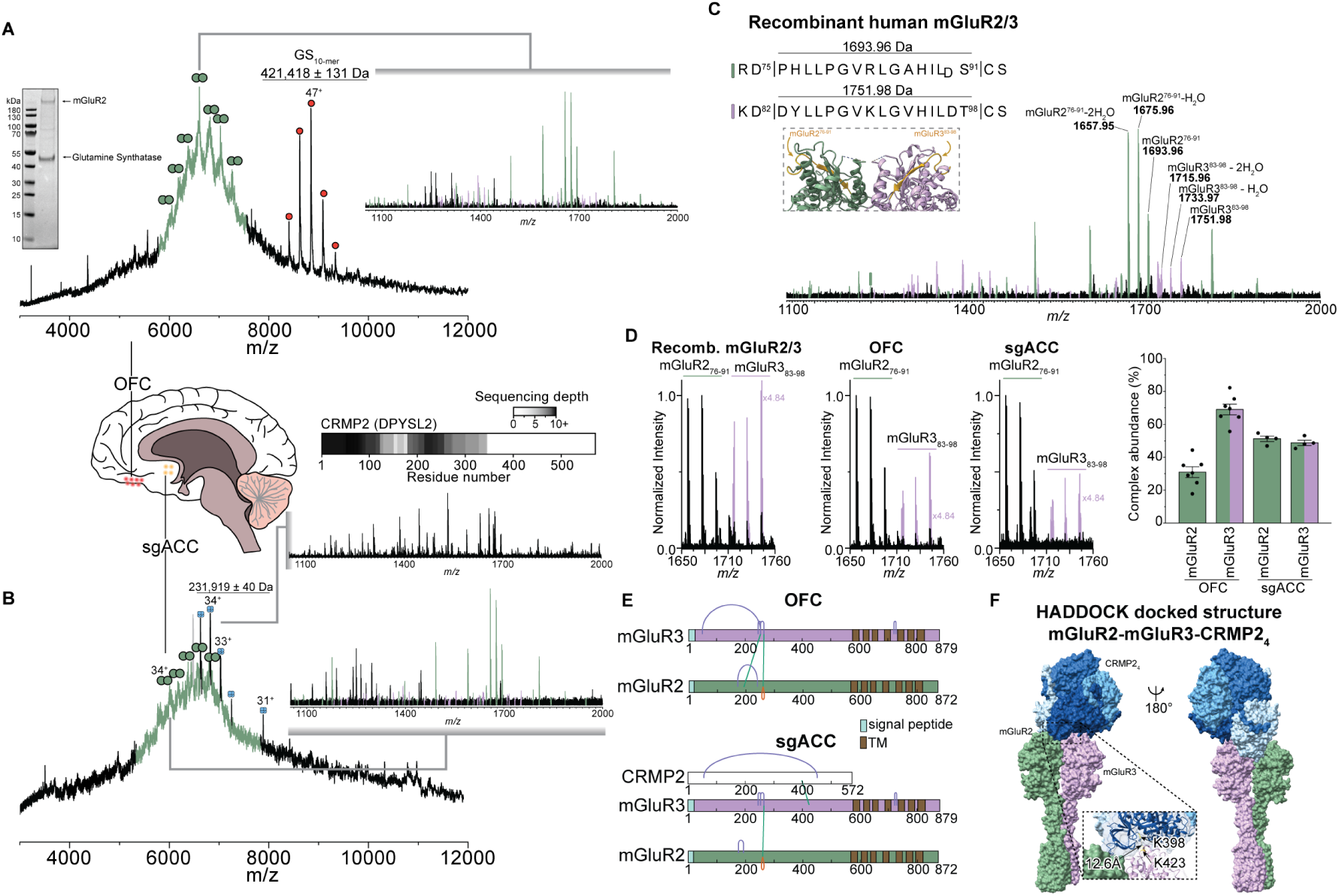
Heterodimeric mGluR2/3 is highly abundant in human OFC and sgACC, and forms regional-specific interactions. (A) native mass spectrum of mGluR2 preparations from OFC (n=8 donors); (left inset) SDS-PAGE gel showing an intense band at ∼42 kDa which corresponds to glutamine synthetase; right inset shows nTDMS spectrum generated by fragmenting the peak indicated. (B) native mass spectrum of mGluR2 prepared from sgACC (n=4) with top inset depicting nTDMS spectra and residue coverage map for the ∼231 kDa peak that was identified as CRMP2. Bottom inset illustrated that nTDMS of the indicated mGluR2-associated dimer peak contains mGluR3 fragments. (C) nTDMS spectrum of recombinant human mGluR2/3; the inset shows the location of mGluR2^76–91^ and mGluR3^83–98^ in the upper lobe of the Venus Fly Trap domain (colored gold). For both mGluR2^482–499^ and mGluR3^495–508^ extensive dehydration is observed, typical for peptides containing acidic residues. (D) quantification of the mGluR2- and mGluR3-associated peaks (inclusive of dehydration products) in recombinant human mGluR2/3 reveals that a 1:1 complex generates fragments with a 4.84:1 intensity ratio (mGluR2^482–499^:mGluR3^495–508^), establishing a calibration baseline. These peaks are present in nTDMS spectra of mGluR2 preparations from OFC and sgACC, albeit at different intensities. After calibration, the bar plots depict the relative mGluR2/2 and mGluR2/3 complex abundances in these anatomically-defined regions. Bar heights denote the mean and error bars the standard error of the mean (n=7 OFC and n=4 sgACC). (E) domain maps depicting the high-confidence crosslinks (FDR<0.05) determined by XL-MS analysis of mGluR2 immunoprecipitates from OFC and sgACC. Crosslinks between mGluR2 and mGluR3 are found in OFC and sgACC; crosslinks between the Venus Fly Trap domain of mGluR3 and CRMP2 are uniquely recovered in sgACC. (F) data-driven protein docking of the mGluR2-mGluR3 heterodimer^57^ extracellular domain with the CRMP2 tetramer,^66^ guided by DSSO crosslinking mass spectrometry. The docking model positions mGluR3 residue K423 in the Venus Flytrap domain within 12 Å of K398 of a CRMP2 protomer, consistent with the experimentally observed crosslink (FDR ≤ 0.05).

We next examined mGluR2 from the sgACC (**Figure 4B**). As in the OFC, dimer charge state peaks were clearly evident, and nTDMS confirmed the presence of both mGluR2 and mGluR3 (**Figure 4B bottom inset**). Unique to sgACC preparations, we also observed a series of sharp peaks corresponding to a complex weighing 231,919 ± 40 Da, interleaved between the mGluR dimer peaks. nTDMS of the 34^+^ charge state peak (**Figure 4B top inset**) produced a fragment spectrum consistent with CRMP2 (gene name DPYSL2), lacking the initiator methionine and bearing an acetylated N-terminus. The CRMP2 protomer mass was 4,225 Da lighter than expected from the sequence, and the lack of C-terminal sequence coverage is consistent with reports of heterogenous proteolytic processing of this disordered region.^68^ CRMP2 is a tetrameric enzyme that is implicated in neuronal development and axonal growth through its regulation of microtubule dynamics, cytoskeletal organization, and vesicular trafficking.^66^ Although canonically localized in the cytosol, CRMP2 has been detected in the extracellular space including in cerebrospinal fluid and conditioned media of neuronal cells (**Figure S6**).^69–71^ It has been suggested that CRMP2 modulates intercellular signalling cascades through unconventional binding mechanisms.^71^ Together, our data for OFC and sgACC confirm that mGluR2/3 heterodimers are present but that GS and CRMP2 interact uniquely with mGluR2/3 in different regions.

Having established the presence of heterodimers we then sought to quantify their levels in the OFC and sgACC using nTDMS. We mined nTDMS spectra of recombinant human mGluR2 and mGluR3 for fragments originating from structurally equivalent regions within the extracellular Venus Fly Trap domain (**Figure 4C and Figure S7**). Intense fragments corresponded to cleavage events at Asp75/Cys92 in mGluR2 and Asp82/Cys99 in mGluR3. Both fragments were present in recombinant human mGluR2/3, albeit at different signal intensities. After accounting for the extensive dehydration products observed for mGluR2^75–92^ and mGluR3^82–99^, the mGluR2:mGluR3 calibration baseline was 4.84:1 (**Figure 4D**). The ∼5-fold disparity in mGluR3 fragments emanating from mGluR2/3 complexes is likely due to sequence- and structure-guided nTDMS fragmentation patterns, which have previously been addressed.^39,41^ Applying this strategy to specimens isolated from human brain, we identified mGluR2^75–92^ and mGluR3^82–99^ in nTDMS spectra from OFC and sgACC (**Figure 4D**). After calibration, high levels of mGluR2/3 heterodimers were observed in both regions: ∼70% in the OFC and ∼50% in the sgACC. While there are high levels of heterodimers in human cortex, there are regional differences in the homodimer and heterodimer ratios, likely linked to a functional outcome.

Given the differential mGluR2/3 interactions in OFC and sgACC, we next examined how co-immunoprecipitating GS and CRMP2 are organized using chemical crosslinking mass spectrometry (XL-MS). In OFC preparations, we recovered DSSO crosslinks across the lower lobes of the mGluR2 and mGluR3 Venus Fly Trap domains, consistent with the formation of stable heterodimers (**Figure 4E**). Crosslinks were not detected between either mGluR2 or mGluR3 subunits and GS, suggesting its co-immunoprecipitation with mGluR2/3 reflects indirect or transient association rather than stable physical contact. sgACC preparations also yielded crosslinks across the Venus Flytrap domains of mGluR2 and mGluR3. A crosslink between CRMP2 and the mGluR3 subunit was identified in the sgACC preparation, indicating a direct and stable interaction (**Figure 4E**).

Guided by the crosslinking results, we performed data-driven protein docking of the CRMP2 tetramer onto the mGluR2/3 heterodimer extracellular domain using HADDOCK;^72,73^ the DSSO crosslink between mGluR3 K423 and CRMP2 K398 was applied as a distance restraint. The top-scoring docking cluster (HADDOCK score −94.2 ± 8.5, z-score −1.8) positioned mGluR3 K423 within 12 Å of CRMP2 K398, well within the maximum DSSO crosslinker span of 35 Å, validating the docking solution against the experimental data (**Figure 4F**). The interface was characterized by a buried surface area of 2134 ± 81 Å^2^ and strong electrostatic complementarity, with CRMP2 contacting the top lobe of the mGluR3 Venus Flytrap domain. Across ten independent docking runs converging to the same solution, the CRMP2 tetramer docked to mGluR3, further suggesting a defined and reproducible binding mode between CRMP2 and the mGluR2/3 heterodimer in the sgACC. These data provide evidence that mGluR2/3 forms unique interactions with their surrounding environments in different neuroanatomically-defined and functionally-distinct regions of the brain.

### Immunoprecipitation of VGluT1 and mGluR2 from a cohort of post-mortem human OFC with depression

Having established the molecular composition and interactions of mGluR complexes in healthy brain sections, we asked whether these assemblies are disrupted in major depressive disorder. We assembled a cohort of fresh-frozen OFC tissue from depressed subjects who died by suicide (DEPR, n=7) age-matched to healthy controls (CON, n=8) (**Figure 5A, Table S3**). Severe depression with completed suicide represents the most clinically homogeneous and biologically extreme end of the depressive spectrum. We did not observe significant differences in VGluT1, mGluR2 and mGluR3 protein expression levels between CON and DEPR groups (**Figure 5B**). We also confirmed that the prolonged post-mortem intervals, typical of biobanked suicide specimens, do not compromise native complex integrity for both VGluT1 and mGluR2 (**Figure S8**). This represents an important validation for the use of post-mortem tissue in nMS.

**Figure 5.**
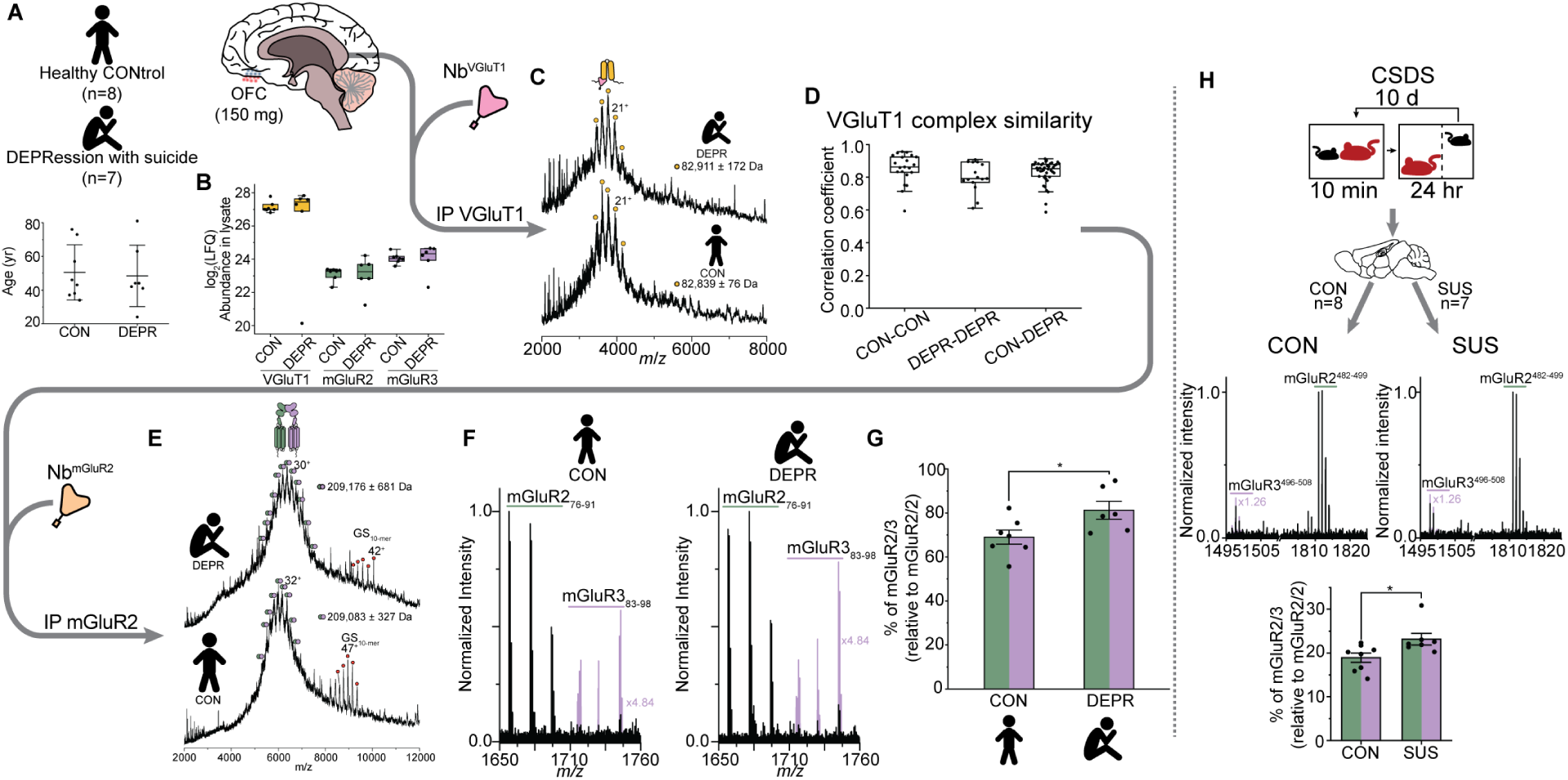
The organization of VGluT1 and mGluR2 in a depression cohort. (A) study design and cohort details. Horizontal lines depicts the mean age and error bars the standard deviation of the mean (CON – 50.5±15.3 yr; DEPR – 48.4±16.9 yr). (B) protein abundances of VGluT1, mGluR2, and mGluR3 in the OFC (n=8 CON, n=7 DEPR). No other mGluRs were observed, suggesting they are present at very low abundances in the OFC. Protein abundances were not different between groups (two-tailed t-test for means, all p-values >0.05). (C) representative native mass spectra of VGluT1 from a CON and DEPR patient; spectra are nearly superimposable indicating VGluT1 complexes are unchanged with psychiatric illness. (D) box plots depicting within- and between-group correlation coefficients determined for VGluT1 native mass spectra. Box widths denote the first and third interquartile range (IQR), horizontal lines the median, and whiskers 1.5x the IQR beyond the box edge. Within group median values (CON-CON: 0.85; DEPR-DEPR: 0.80) and between-group values (CON-DEPR: 0.83) are similar, indicating that there are minimal differences in protein interactions for VGluT1 in DEPR relative to CON. (E) representative native mass spectra of mGluR2 immunoprecipitated from CON and DEPR patients; dimer masses are similar, and GS_10-mer_ persists in both groups. There is a slight broadening of the dimer peak charge state distributions, suggestive of compositional heterogeneity. (F) nTDMS of a single dimer peak from CON and DEPR patients, with regions for mGluR3 scaled by 4.84x for quantification. (G) bar plot quantifying the relative abundance of mGluR2/3 in CON and DEPR subjects (two-tailed t-test for means, n=7 CON, n=6 DEPR, p=0.03). Bar heights depict the mean and error bars the standard error of the mean. Only high quality nTDMS datasets were used for quantification (methods). (H) mGluR2 was immunoprecipitated from mice subjected to CSDS and preparations were analyzed by nTDMS, revealing a modest but significant positive correlation between % mGluR2/3 and SUS vs CON (n=8 CON and n=7 SUS, two-tailed t-test for means, p = 0.04). Bar heights depict the mean and error bars the standard error of the mean.

Considering MS detail per-patient, we find that for VGluT1 complex masses, lipid and ligand binding patterns, within- and between CON and DEPR group yield correlation coefficients, that are indistinguishable (**Figures 5C-D**, median r > 0.8). Moreover, nTDMS revealed no differences in PTM status or sequence variation (**Figure S9**). VGluT1 complexes are therefore remarkably stable across the disease boundary. The molecular organization of VGluT1, and by extension, the machinery of presynaptic glutamate loading, therefore appears to be preserved in the OFC of patients with severe depression. This would imply that the primary defect in this disorder does not lie at the level of vesicular glutamate packaging.

Turning our attention to the mGluR2 immunoprecipitates mGluR2 dimer masses were similar across groups and copurification of decameric glutamine synthetase persisted in both CON and DEPR (**Figure 5E**). nTDMS quantification however revealed a striking shift in homodimer and heterodimer ratios. Fragments diagnostic for mGluR3 were consistently elevated in DEPR relative to CON (**Figure 5F**). The mean proportion of mGluR2/3 heterodimers was significantly increased to ∼80% in DEPR, an elevation of ∼15 percentage points from baseline levels in healthy controls (*viz.* a ∼21% increase) (**Figure 5G**). This finding indicates that depression is associated not with a change in total mGluR2 expression, but with a specific organization of receptor subunit composition, favoring heterodimeric mGluR2/3 over mGluR2 homodimers in the OFC.

To cross-validate this finding of elevated mGluR2/3 levels in a controlled experimental system, we immunoprecipitated mGluR2 from mice subjected to chronic social defeat stress (CSDS), a well-established paradigm for inducing depressive-like behaviors in rodents.^74–77^ mGluR2/3 levels although lower in mice than in humans (see above) were elevated in SUS animals (∼19% in CON and ∼24% in SUS, **Figure 5H**). Taken together, the convergence of this finding across species (despite absolute differences in baseline mGluR2/3 levels) strengthens our assertion that mGluR2/3 heterodimers are a conserved molecular correlate of depressive states.

## Discussion

We devised a fast immunoprecipitation strategy to capture multiple receptors from individual regions (∼30 – 450 mg) of animal and human brain tissue. By coupling this capability with a multitude of downstream MS studies, we demonstrated the molecular organization of VGluT1 and mGluR2 from a single mouse brain. Our findings revealed that endogenous VGluT1 was glycosylated at a single N-glycan site, and formed extensive interactions with phospholipids. From this same mouse brain, immunoprecipitated mGluR2 was heavily glycosylated with each N-glycan site showing limited compositional diversity. Together, these observations are in good agreement with recent glycoproteomic and imaging studies indicating that the majority of N-glycans in the brain are compositionally-restricted to high mannose and short hybrid/complex type.^78,79^ Endogenous mGluR2 also formed extensive interactions with neutral and anionic phospholipids from the membrane, which underly the broad heterogeneity in dimer molecular weight distribution. Lastly, by comparing nTDMS datasets collected from endogenous murine mGluR2 with recombinant murine mGluR2/2, mGluR3/3 and mGluR2/3, we identified clear evidence for low populations (∼ 20%) of endogenous mGluR2/3 heterodimers, from individual mouse brain specimens.

This exciting finding led us to explore the organization of heterodimers across neuroanatomically and functionally distinct regions of the mouse brain. Between whole brain and cerebral cortex, mGluR2/3 heterodimers were detected at similar abundance levels (∼20%), while immunoprecipitations from hippocampus and cerebellum yielded comparatively lower levels (∼10%) (**Figure 6A**). The approximately two-fold enrichment in cortex relative to hippocampus and cerebellum may reflect a greater demand for heterodimer-specific properties at cortical synapses, where glutamatergic signalling is particularly dense.

**Figure 6.**
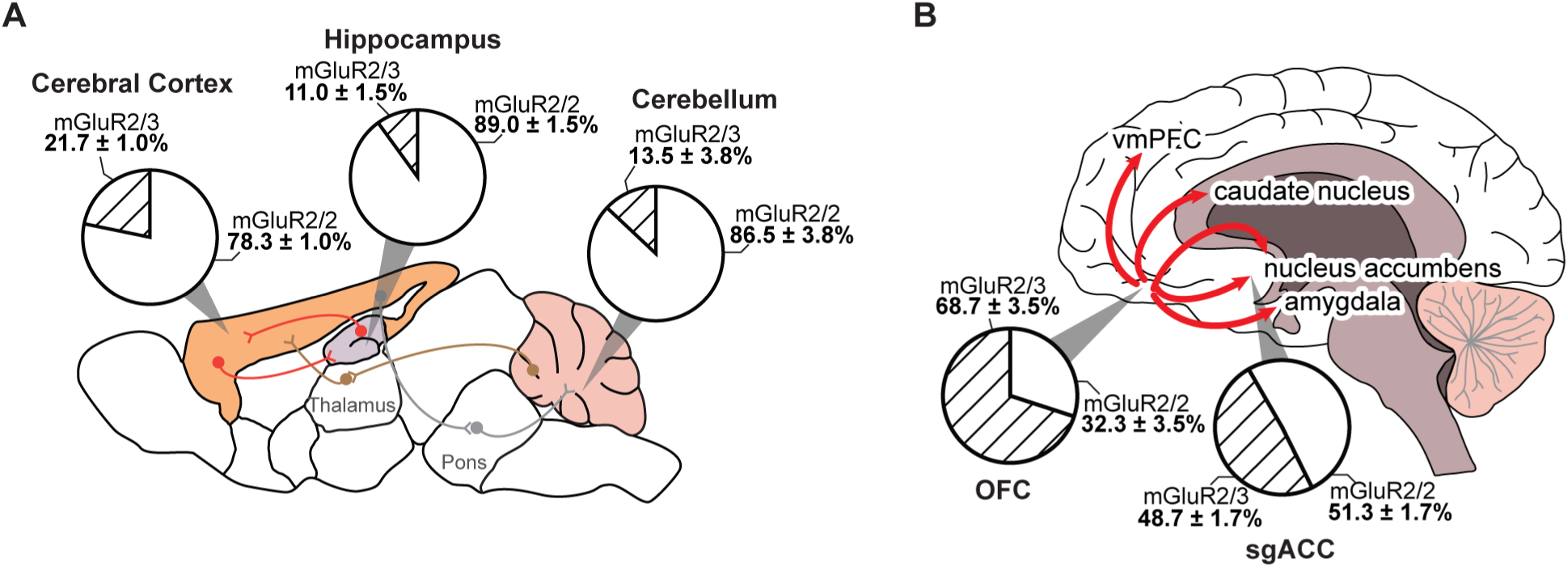
Regional distribution of mGluR2/3 heterodimer levels in mouse and human brain. (A) Proportion of mGluR2/3 heterodimers detected across mouse brain regions sampled in this study. Symbols indicate synaptic connectivity pathways between regions. (B) Heterodimer proportions in two human prefrontal cortical regions. The OFC exhibits a higher heterodimer burden (∼68% mGluR2/3) relative to the sgACC (∼48% mGluR2/3). The arrow indicates the directional projection from OFC to subcortical and cortical targets including the nucleus accumbens, amygdala, caudate nucleus, vmPFC, and sgACC. mGluR2/2, homodimer; mGluR2/3, heterodimer.

High levels of mGluR2/3 heterodimers (∼70%) were detected in the human OFC, a highly connected region that projects to subcortical and cortical targets including the nucleus accumbens, caudate nucleus, amygdala, vmPFC, and sgACC (**Figure 6B**).^59,60,80^ Through lesion studies in non-human primates^1,58^ and neuroimaging in humans,^61^ the OFC is linked to incentive salience, decision-making, anticipatory processing, and mood regulation.^1,59,60^ By contrast, the sgACC, a region critically implicated in emotional dysregulation, autonomic control, and vulnerability to mood disorders,^1,62^ exhibited comparatively lower heterodimer levels (∼50%). The OFC-to-sgACC projections are noteworthy; as part of a hierarchical prefrontal affective circuit (**Figure 6B**),^1,81,82^ the heterodimer gradient observed between these regions raises the possibility that mGluR2/3 assembly is differentially regulated across successive nodes of this pathway. Moreover, the identification of CRMP2 as a binding partner of the mGluR2/3 heterodimer in the sgACC suggests a previously unrecognized protein complex. Although CRMP2 is canonically localized to the cytosol where it is involved in microtubule organization, axonal growth, and cytoskeletal regulation,^66,68^ extracellular detection^69–71^ indicates that an extracellular population is positioned to interact with the mGluR2/3 Venus Flytrap domain.

These structural and regional observations invite several mechanistic hypotheses. The tetrameric architecture of CRMP2 could confer an extracellular scaffolding function, simultaneously engaging multiple mGluR2/3 heterodimers to organize perisynaptic receptor clusters. The positioning of CRMP2 at the upper lobe of the mGluR3 Venus Flytrap domain raises the possibility that it acts as an allosteric modulator of glutamate-induced receptor activation, analogous to the behavior of allosteric mGluR2 nanobodies.^49^ The regional specificity of this interaction to the sgACC is particularly noteworthy given the established roles of mGluR2/3 signaling in corticolimbic glutamate regulation in psychiatric disorders.^83^

The finding that mGluR2/3 heterodimer abundance is selectively elevated in the OFC of individuals with severe depression (without changes in total receptor expression or VGluT1 complex organization) points to reorganization of synaptic receptor architecture as a molecular correlate of disease. This is conceptually important: it suggests that a relevant correlate is not how much receptor is present, but how it is assembled. Given that mGluR2 and mGluR3 homodimers and mGluR2/3 heterodimers differ in their signalling properties, G protein coupling efficiency, and sensitivity to allosteric modulators,^10,13–15,19,20,23,24,84–86^ a shift in their relative abundance could alter the gain of presynaptic inhibitory control over glutamate release in OFC circuits implicated in mood and decision-making. The convergence of this finding in the CSDS mouse model strengthens the case that heterodimer upregulation is not an epiphenomenon of post-mortem confounds or cohort-specific factors, but a conserved biological correlate to depressive states across species. Notably, while the absolute heterodimer levels differ substantially between human OFC and mouse, the directional shift in disease conditions is preserved. This suggests that the dynamic regulation of the homo/heterodimer balance may represent a physiologically and pathologically relevant correlate. As such these findings imply that selectively modulating these assemblies may offer greater precision than non-selective mGluR2/3 modulators.

Considering our findings in a broader context, previous studies have reported that mGluR subtypes can form heterodimers^21^ across the cortex^22^ with atomic structures from recombinant systems reported^19,57^. Evidence for mGluR2/4 heterodimers in cells and *in vivo* have been documented^14,15,22^. However, direct observation of mGluR2/3 heterodimers from human brain tissue, together with associated lipids and non-canonical binding partners, has remained elusive. We provide a strategy for dissecting endogenous synaptic complexes from brain sections from humans and animals with molecular resolution, demonstrating existence of mGluR2/3 heterodimers in healthy mice (∼18%) and humans (∼65%). This stark species difference, together with the unique pharmacological profile of mGluR heterodimers^13,15,87^, may help explain the tempered success in translating mGluR-specific modulators from rodent studies to the clinic^88^. More broadly, the demonstration that receptor assembly state, rather than expression level, tracks with disease severity reframes how the field approaches target engagement and biomarker development for glutamatergic therapies. We anticipate that the molecular autopsy framework developed here, enabling detailed analyses of receptor assembly states, post-translational modifications, associated lipids, and other binding partners from small amounts of individual tissue sections, will be applicable across multiple receptor families and brain disorders. This approach promises a mechanistically grounded, molecular-level understanding of how the organizations of endogenous receptor complexes shapes neuronal computation, behaviour, and vulnerability to disease, with implications across the full spectrum of brain disorders - from psychiatric conditions to pain, consciousness, and the neurobiology of mind-altering drugs.

### Limitations of the study

Molecular studies require the removal of receptors from their native cell membranes, which risks the loss of interactions that depend on intact membrane context. Furthermore, tissue lysis precludes linking mGluR heterodimer levels to specific cell types, and manually dissected regions such as OFC and sgACC will inevitably contain cytoarchitecturally heterogeneous tissue. mGluRs are disulfide-linked dimers, which minimises the risk that altered heterodimer levels reflect lysis-induced receptor remodelling rather than native assembly states. Current hardware constraints restrict non-covalent lipid detection to positively charged species, leaving anionic phospholipids (including phosphoinositides and phosphatidylserine, which are functionally relevant to GPCR signaling) undetected in our current nTDMS instrument arrangement. This can be addressed in parallel using nanoflow lipidomics on immunoprecipitates as done in this study. While the focus of this work is on method development, we note that cohort sizes are modest. Nonetheless, we observe reproducible compositional heterogeneity attributable to mGluR2/3 heterodimer levels across both human and mouse cohorts. The human cohort is restricted to individuals with severe depression who died by suicide, limiting generalizability to non-suicidal depression, and potential confounders including medication history, substance use, agonal state, and sex composition cannot be fully excluded. Post-mortem studies are inherently cross-sectional, which limits our ability to establish whether heterodimer reorganisation is a cause or consequence of depressive pathology; longitudinal or experimental paradigms will be needed to resolve this question.

## Acknowledgements

T.J.E. acknowledges Brian Shoichet (UCSF) for invaluable discussions on co-translational assembly. Bo Huang (UCSF) and James Wells (UCSF) are thanked for discussions on strategies to detect heterodimers *in vivo*. The authors are grateful to Diego A. Pizzagalli (McLean Hospital/Harvard Medical School and UC, Irvine), Helen S. Mayberg (Icahn School of Medicine at Mount Sinai), Jonathan Flint (UCLA), and Michelle G. Craske (UCLA) for their insightful comments over the course of this study. John E.P. Syka, Joshua D. Hinkle, Rafael D. Melani, and Christopher Mullen (Thermo Fisher Scientific, San Jose, CA) are acknowledged for their critical support with the development of the IRMPD modification to the Orbitrap Ascend instrument. Alexander Makarov (Thermo Fisher Scientific Bremen, Germany) is thanked for his guidance with tuning the Orbitrap UHMR instrument for high molecular weight protein complexes. T.J.E., T.R., and O.B.R. thank Randy Bruno (Department of Physiology, Anatomy, and Genetics, University of Oxford) for advice on murine tissue dissections. T.J.E., C.A.L., and C.V.R. thank Nicholas M. Riley and Carolyn R. Bertozzi (Stanford University) for providing access to lab space during a critical stage in instrument development. We thank Robert Hedley and Vasiliki Tsioligka for providing technical assistance with fluorescence-assisted cell sorting at The Don Mason Facility of Flow Cytometry, Sir William Dunn School of Pathology, University of Oxford.

## Data availability

All mass spectrometry data have been deposited to the ProteomeXchange Consortium via the PRIDE^89^ partner repository with the dataset identifier PXDXXXXXX. Plasmids and cell lines generated in this study will be made available upon reasonable request to the corresponding authors.

## Author Contributions

T.J.E., C.A.L., and C.V.R. conceived and designed the study. T.J.E. designed and generated expression constructs with assistance from C.A.L., S.A.S.L., and F.I.B. T.J.E., C.A.L., J.L.B., S.A.S.L., O.B.R., T.R., and F.I.B. generated cell lines, carried out cell culture, purified recombinant proteins, and expressed and purified Nbs. T.J.E. performed immunoprecipitations. C.A.L., T.J.E., and J.L.B. collected and analyzed nMS and nTDMS data. T.J.E., S.A.S.L., O.B.R., T.R., S.A.B., and W.S. performed and analyzed proteomics and glycoproteomics. T.J.E. and S. A. B. carried out crosslinking-mass spectrometry. C.A.L. and D.W. carried out lipidomic studies. H.S. assisted with cryoEM structural evaluation; K.L.C., L.F.P., E.M.P and S.J.R. carried out chronic social defeat stress procedures. T.J.E. and O.B.R. dissected murine tissue. J.W.M. gave input to the study design and data interpretation. T.J.E. and C.V.R. wrote the manuscript with input from all authors.

## Methods

### Statistical approaches

No statistical approach was used to determine sampling size, and authors were not blinded to experimental conditions. Statistical significance was determined using a threshold of p-value < 0.05. Bonferroni-Holm corrected p-values were used as indicated to account for multiple hypothesis testing.

### Expression of nanobodies

Codon-optimised (*E. coli* expression) synthetic genes encoding nanobody (Nb) DN07 targeting mGluR2^49^ (WO2016001417A1) and Nb DN09 targeting VGluT1^48^ were synthesised by Integrated DNA technologies. The constructs contained an N-terminal PelB leader sequence and a C-terminal FLAG-tag. Genes were cloned into an in-house modified pet15b vector using In-Fusion cloning (Takara Bioscience). The plasmid sequences were verified by Sanger and nanopore sequencing.

Nb^VGluT1^ and Nb^mGluR2^ were overexpressed in *E. coli* BL21(DE3) cells in a similar manner as previously described ^18^. Briefly, BL21(DE3) cells were transformed with the assembled plasmids by heat shock and grown overnight on LB agar plates supplemented with 100 µg mL^-1^ ampicillin. The following day, 5–10 colonies were selected to inoculate 500 mL of LB broth (supplemented with 100 µg mL^-1^ ampicillin and 1% glucose). ∼10 mL of this overnight culture was used to inoculate 1 L of LB broth supplemented with 100 µg mL^-1^ ampicillin and 1% glucose. Protein expression was induced by adding 1 mM IPTG to cultures in mid-log phase (OD_600nm_ ≈ 0.4–0.8). Nbs were allowed to express overnight at 28°C. The following day, cells were pelleted by centrifugation (5000×*g*, 4°C) and resuspended in TES buffer (20 mM Tris [pH 8.0 at 4°C], 500 µM EDTA, 500 mM sucrose) for ∼1 hr at 4°C. Outer membranes were ruptured *via* osmotic shock by adding an equal volume of 0.25X TES buffer to release the periplasmic fraction (1 hr, 4°C).

The periplasmic fraction was separated from cell debris via centrifugation (20000×*g,* 1 hr, 4°C) and passed over Ni-NTA resin pre-equilibrated with HNG buffer (50 mM HEPES pH 8.0, 150 mM NaCl, 5% glycerol). The column was washed with >10 column volumes of HNG buffer supplemented with 20 mM imidazole, and proteins were eluted using HNG buffer supplemented with 500 mM imidazole. Imidazole was subsequently removed via overnight dialysis against HNG buffer with a 3 kDa MWCO dialysis cassette (Pierce). Nbs were concentrated to ∼1–2 mg mL^-1^, aliquoted, snap-frozen in liquid nitrogen (LN_2_), and stored at −80°C.

### Cells and cell culture

HEK293S GNTI^-/-^ (ATCC-CRL-3022) was obtained from LGC standards. Lenti-X cells were obtained from Takara. Cell lines were used without authentication and routinely tested negative for mycoplasma. Cells were cultured as adherent monolayers in 1X DMEM:F12 (Life Technologies, Thermo Fisher Scientific) supplemented with 10% fetal bovine serum (FBS) (Life Technologies, Thermo Fisher Scientific) and 1X non-essential amino acids (Life Technologies, Thermo Fisher Scientific).

### Expression and purification of VGluT1

A codon-optimised synthetic gene encoding full length human VGluT1 with a C-terminal FLAG-tag and Twin-Strep-tag II was synthesised by Integrated DNA technologies. The resulting gene was cloned into pcDNA 3.1(+) between the NotI and ApaI restriction sites. The plasmid was electroporated into HEK293 GNTI^-/-^ cells using the manufacturer’s recommended settings (Invitrogen). Cells were allowed to grow for at least 21 days with periodic passaging in 1X DMEM/F12 supplemented with 10% FBS, 1X non-essential amino acids, and 400 µg mL^-1^ G418 (Life Technologies) at 37°C and 5% CO_2_ in a humidified incubator. Monoclonal lines were used for overexpression after adaptation to suspension growth as described previously^90^, except that 1X FreeStyle media (supplemented with 1% FBS) was used in place of 1X DMEM. Cells were seeded into ∼400 mL of suspension media and cultured at 37°C in a humidified incubator (8% CO_2_ and 120 rpm) until reaching a density of ∼3.0 × 10^6^ cells mL^-1^. Cells were then pelleted at 500×*g* for 10 min, washed once in ∼30 mL of phosphate buffered saline, pelleted again at 500×*g* (10 min), snap frozen in LN_2_, and stored at −80°C. Cells were resuspended in HNG buffer supplemented with cOmplete protease inhibitor cocktail (Roche) and lysed with two passages through a microfluidiser (12,000 psi). Unbroken cells and debris were pelleted at 20000×*g* (4°C, 20 min) prior to the isolation of cell membranes using ultracentrifugation (180000×*g*, 4°C, 1 hr). Membranes were solubilised in HNG buffer supplemented with 1% (*w*/*v*) n-dodecyl-β-D-maltoside (DDM)/0.1% cholesteryl hemisuccinate (CHS) for 1 hr at 4°C with rotation. Insoluble material was pelleted (20000×*g*, 4°C, 20 min), and the solubilised membranes were incubated with anti-DYKDDDK magnetic agarose resin (Life Technologies, Thermo Fisher Scientific) for 1 hr. The resin was washed extensively with HNG buffer supplemented with 0.017%/0.0017% DDM/CHS prior to elution using 1 mg mL^-1^ DYKDDDDK peptide (Genscript). VGluT1 was concentrated to ∼5 µM, aliquoted, snap frozen in LN_2_, and stored at −80°C.

### Expression and purification of mGluR2, mGluR3, and mGluR2/3

Codon-optimised (human or mouse expression) synthetic DNAs encoding full-length mGluR2 and GRM2 were purchased from IDT DNA technologies. For affinity purification, human and mouse mGluR2 harbored a C-terminal TEV-AVI-(GSSS)_2_-StrepII tag and mGluR3 harbored a C-terminal TEV-FLAG-(GSSS)_2_-StrepII tail. Mouse mGluR3 harbored a C-terminal TEV-(GSSS)_2_-StrepII tail. These DNA constructs were separately cloned into lentiviral transfer plasmids with an eGFP reporter gene (a kind gift from J. Elegheert and R. Aricescu, Addgene plasmid #113884) between EcoRI and XhoI restriction sites. Lentiviruses packing mGluR2-IRES-eGFP and mGluR3-IRES-eGFP were generated as described ^91^ using pMD2.G lentiviral envelope plasmid (Addgene, cat. no. 12259) and psPAX2 lentiviral packaging plasmid (Addgene, cat. no. 12260), which were kind gifts from Didier Trono’s laboratory. Supernatants (18 mL) containing high-titer lentiviral particles were used to infect HEK293 GNTI^-/-^ cells as described. Cells positive for eGFP were separated from non-transduced cells using fluorescence-activated cell sorting (FACS) at the William Dunn School of Pathology, University of Oxford. Polyclonal lines were adapted to suspension growth in 1X FreeStyle supplemented with 1% FBS. Upon reaching a density of ∼3.0–4.0 × 10^6^ cell mL^-1^, 5 mM sodium butyrate was added to boost protein expression for an additional 24 hr under the same growth conditions. Membranes from ∼1L of cells were solubilized in HNG buffer supplemented with 1% DDM)/0.1% CHS for 1 hr at 4°C with rotation. Insoluble material was pelleted (20000×*g*, 4°C, 20 min), and the solubilised membranes were incubated with anti-DYKDDDK magnetic agarose resin (Life Technologies, Thermo Fisher Scientific) for 1 hr. To delipidate, the immobilise proteins were twice incubated with HNG buffer supplemented with 2% DDM/0.2% CHS for ∼3 hrs each time. The delipidated proteins were washed extensively with HNG buffer supplemented with 0.017%/0.0017% DDM/CHS prior to elution using 1 mg mL^-1^ DYKDDDDK peptide (Genscript). mGluR2/2 and mGluR3/3 were concentrated to ∼10 µM (dimer concentration), aliquoted, snap frozen in LN_2_, and stored at −80°C.

To generate mGluR2/3 heteromeric complexes, mGluR2 was cloned into a lentiviral transfer plasmid with an mRuby reporter (a kind gift from J. Elegheert and R. Aricescu, Addgene plasmid #113885). A heterogenous distribution of lentiviral particles was generated in a single Lenti-X cell population by co-transfecting mGluR2-IRES-mRuby and mGluR3-IRES-eGFP along with psPAX2 and pMD2.G plasmids. The supernatant was used to infect HEK293 GNTI^-/-^ cells, and cells positive for both mRuby and eGFP were sorted via compensated FACS as described^91^. Cells were cultured for ∼2 weeks at 37°C (5% CO_2_ in a humidified incubator) in DMEM:F12 supplemented with 10% FBS and 1x non-essential amino acids. mGluR2/3 expressing cells were adapted to suspension growth in ∼1 L of FreeStyle media (supplemented with 1% FBS) at a starting density of ∼0.5 x 10^6^ cells mL^-1^ and grown at 37°C (8% CO_2_, 120 rpm). At a density of ∼3.0–4.0 × 10^6^ cell mL^-1^, 5 mM sodium butyrate was added to boost protein expression for an additional 24 hr under the same growth conditions. Membranes from 1 L of culture were isolated as described for VGluT1.

On the day of purification, solubilised membranes from ∼1L of culture were incubated with anti-DYKDDDK magnetic agarose (Life Technologies, Thermo Fisher Scientific) resin for 1 hr prior to separation from the lysate and subjected to delipidation as described above. The delipidated mGluR2/3 complexes were extensively washed with HNG buffer (supplemented with 0.017%/0.0017% DDM/CHS). FLAG-tagged mGluR2/2 and mGluR2/3 containing complexes were eluted twice (1 hr per elution) with 1 mg mL^-1^ synthetic FLAG peptide (Genscript). The resulting eluate was incubated with MagStrep “type3” XT beads pre-equilibrated with HNG buffer supplemented with 0.017%/0.0017% DDM/CHS for at least 1 hr. The resin was extensively washed with HNG buffer supplemented with 0.017%/0.0017% DDM/CHS, prior to elution with HNG supplemented with 0.017%/0.0017% DDM/CHS and 50 mM biotin. The resulting mGluR2/3 containing complexes were concentrated to ∼5 µM (dimer), aliquoted, snap frozen in LN_2_, and stored at −80°C.

### Mouse brains used for Fig 1 and 2

The Nb immunoprecipitation approach was developed using reclaimed brain tissue harvested from adult C57BL6 mice, provided by BioIVT (England, UK). For whole brain immunoprecipitations (**Figure 1 and 2**), cryogenically-frozen whole brains were immersed in a liquid nitrogen bath and crushed using a mortar and pestle. The powder was resuspended in lysis buffer and processed as described above. For regional studies, whole C57BL6J mouse brains were thawed in ice-cold PBS supplemented with protease inhibitors (Roche). Regions were rapidly dissected using the Allen Brain Atlas as a reference and snap-frozen prior to processing. Regions were resuspended in ice-cold lysis buffer and homogenizing by gentle pipetting using a P200 pipette. Cells were further lysed by passing homogenates through a 27G needle prior to lysis and solubilization (see below). All steps were carried out at 4 °C.

### Mice for chronic social defeat stress (CSDS)

Chronic social defeat stress experiments used 7-week-old male C57BL/6J mice (stock no. 000664) purchased from The Jackson Laboratory. Four to six-month old retired male CD-1 breeders were used as aggressors (purchased from Charles River Laboratories, Crl:CD1[ICR]). To select for aggressive CD-1 mice for use in defeat experiments, we selected aggressive behaviors towards C57BL/6J mice prior to their use. Mice were maintained on a 12 hr light:dark cycle (7:00 am lights on, 7:00 pm lights off) with access to food and water *ad libitum*. Mice were acclimated to the testing room for >1 hr prior to behavioral assays. All procedures were performed in accordance with the National Institutes of Health Guide for care and Use of Laboratory Animals and the Icahn School of Medicine at Mount Sinai Institutional Animal Care and Use Committee.

### CSDS and social interaction (SI) test

CSDS was carried out as previously described^75,92^. Briefly, retired male breeder CD-1 mice were aggressors and were screened for consistent aggressive behavior (at least one aggressive encounter per minute) during a pre-experiment screening period. 8-week-old males C57BL/6J were experimental mice. Two days before the start of the defeat, the CD-1 mice were housed on one side of a cage partitioned by perforated plexiglass. Defeat was conducted over 10 consecutive days. Experimental mice were subjected to direct physical interaction with a CD-1 mouse for 10 min per day, with experimental mice exposed to a new aggressor for each of the 10 days. For the rest of the day, the experimental mice were placed on the other side of the plexiglass divider to allow for sensory contact. Wounding was graded on a previously established scoring system^76^ with mice exceeding endpoint levels of wounding removed from the study. In this study and consistent with others, there were no correlations found between wound score and stress-susceptibility^74^.

SI ratios were determined one day after the final defeat session (day 11) and carried out under red light conditions. Mice were habituated to a behavioral suite for 1 hr before behavioral testing, then placed into a plexiglass open field arena (42 cm x 42 cm x 42 cm, Nationwide Plastics) with a mesh enclosure on one end. To determine SI ratios, experimental mice were allowed to freely explore the arena for 2.5 min during which time locomotor activity was tracked and recorded using a Noldus Ethovision System (Noldus Information Technology, version 11.0). After 2.5 min, the experimental mouse was removed, and the arena was cleaned with 70% ethanol. An unfamiliar CD-1 male mouse (novel social target) was then placed in the mesh enclosure, after which time the experimental mouse was returned. The locomotor activity of the experimental mouse was subsequently tracked for an additional 2.5 min. SI ratio was calculated by determining the ratio between the time the experimental mouse spent in the vicinity of the enclosure when a target mouse was present versus absent. Experimental mice with SI ratios ≥ 1 are deemed resilient, and those with an SI ratio < 1 are susceptible to CSDS. Mice susceptible to CSDS (n=7) and control C57BL/6J mice (n=8) were sacrificed via decapitation. Brains were immediately removed and snap frozen in dry ice-cooled isopentane and stored at - 80°C until processing for immunoprecipitation.

### Human Brain Tissue

Brain tissue from the sgACC/Broadmann area 25 was provided by the University of Oxford Brain Bank. Inclusion criteria: geriatric depression score <10, PMI < 30 hrs. Tissue was selected and dissected by Marie Hamard and Prof. Laura Parkkinen (Nuffield Department of Clinical Neuroscience, University of Oxford).

Brain tissue from the OFC/Brodmann area 11 was provided by the Edinburgh Brain Bank, as part of the UK Brain Bank under request TR95/21. Inclusion criteria: No neurodegenerative remarks, death ruled a suicide, PMI <120 hr, documented history of depression. Eight age- and sex-matched controls were procured. The tissue was selected and provided by Prof. Colin Smith (Department of Neuropathology, University of Edinburgh) as part of the MRC Brain Bank Network and is subject to the ethical approvals governed by the U.K. Medical Research Council.

### Nb-based immunoprecipitation of VGluT1 and mGluR2 from brain tissue

Frozen powder or tissue sections from 30–500 mg of mouse or human brain tissue was lysed in HNG buffer supplemented with 1% DDM/0.1% CHS and a protease/phosphatase inhibitor cocktail (Thermo Fisher Scientific) using a tight-fitting Potter-Elvehjem tissue grinder (5–10 strokes, minimal rotation). Cells and membranes were allowed to solubilise with rotation for 1 hr prior to removal of insoluble debris and unbroken cells via centrifugation (15000×*g*, 20 min, 4°C). A further 100 μL of lysate was reserved for proteomic analysis, snap frozen in LN_2_, and stored at −80°C until use.

During tissue lysis, FLAG-tagged Nb was coupled to anti-DYKDDDDK magnetic agarose resin with gentle rotation for 1 hr. The resin was washed with ∼30 column volumes of HNG buffer supplemented with 0.017%/0.0017% DDM/CHS to remove uncoupled Nb. The resulting resin was combined with solubilized brain lysate and allowed to incubate at 4°C ∼4 – 5 hr (to overnight) with gentle rotation. The following day, the resin was separated from the brain lysates, and then washed extensively using HNG buffer supplemented with 0.017%/0.0017% DDM/CHS. Nbs and Nb-target protein complexes were eluted from the resin using 1 mg mL^-1^ DYKDDDDK peptide in 1 mL HNG (supplemented with 0.017%/0.0017% DDM/CHS). Elution was carried out for 1 hr at 4°C with gentle rotation. The elution was repeated for a total of two elution steps which were pooled. 100 μL of the eluate was reserved for proteomic analysis (snap frozen in LN_2_ and stored at −80°C) and 12 μL of eluate was also reserved for lipidomics (snap frozen in LN_2_ and stored at −80°C). The remaining ∼900 µL was concentrated to ∼1 µM target protein using a 100 kDa MWCO filter (Amicon); excess Nb was removed by ultrafiltration if necessary.

VGluT1 was captured first using Nb^VGluT1^, and mGluR2 captured the following day using Nb^mGluR2^ from the exact same lysate. We tested the impact of reversing the purification order, and did not observe any substantial differences in recovery (not shown).

### Liquid chromatography-MS (LC-MS) based proteomic analysis of tissue lysates

Lysates (∼100 μg) were thawed on ice and unfolded in 3.0 M urea (in 100 mM ammonium bicarbonate, pH 7.5, Sigma-Aldrich) for 1 hr. Disulfide bonds were reduced with 1 mM tris 2-carboxyethyl phosphine hydrochloride (Sigma-Aldrich) at 56°C for 1 hr. The reduced cysteine residues were then alkylated with iodoacetamide (2 mM, Sigma-Aldrich) in the dark for 1 hr at room temperature. The urea solution was diluted to 1 M with 100 mM ammonium bicarbonate, and trypsin was added at a 1:50 *w*/*w* ratio (modified sequencing grade trypsin, Promega). Proteins were digested at 37°C overnight. The following day, neat LC-MS grade formic acid (Fisher Scientific) was added to a final concentration of 5% (*v*/*v*), and peptides were subsequently desalted using C18 Spin Tips (100 µL, Pierce). Desalted peptides were dried in a vacuum concentrator and resolubilised in 50 μL of buffer A (0.1% formic acid, LC-MS grade H_2_O, Fisher Scientific). Peptides were loaded onto a C18 trap column (Acclaim PepMap 100, 75 μm × 2 cm, 3 μm; Thermo Fisher Scientific) using an Dionex Ultimate 3000 UHPLC (Thermo Fisher Scientific) at a flow rate of 20 μL min^-1^. Trapped peptides were washed with 60 µL of buffer A, then separated over a 50 cm C18 nano HPLC column (Acclaim PepMap 100,75 μm × 50 cm, 2 μm, Thermo Fisher Scientific) using a 100 min linear gradient of 5–40% buffer B (0.1% formic acid, 80% acetonitrile, 20% H_2_O, Fisher Scientific) at a flow rate of 300 nL min^-1^ (40°C). The eluant from the column was electrosprayed in positive mode into an Orbitrap Eclipse Tribrid mass spectrometer (Thermo Fisher Scientific) operated in a data-dependent acquisition mode (3 s cycle time). Precursor peptides were analyzed in the Orbitrap analyzer (resolution of 120000 @ 200 *m*/*z*, scan range: 300–2000 *m*/*z*, 100% AGC). Precursor ions (charge states 2–5) above an intensity threshold of 5.0 × 10^4^ were selected with the quadrupole using a 0.7 *m*/*z* selection window and fragmented using HCD (30% normalised energy). MS^2^ spectra of dissociated peptide ions were collected in the Orbitrap (resolution of 30000 @ 200 *m*/*z*, 100% AGC). Additional MS^2^ scans of precursor ions (10 ppm tolerance) were dynamically excluded after the initial scan for 30 s.

### LC-MS based proteomic analysis of immunoprecipitates

Eluates from the IP of VGluT1 and mGluR2 (100 μL) were thawed on ice and processed as described above for lysates. Briefly, proteins were unfolded in 3.0 M urea (in 100 mM ammonium bicarbonate, pH 7.5, Sigma-Aldrich) for 1 hr, reduced in 1 mM tris 2-carboxyethyl phosphine hydrochloride (Sigma-Aldrich) at 56°C for 1 hr, and alkylated with iodoacetamide (2 mM, Sigma-Aldrich) in the dark for 1 hr at room temperature. The urea solution was subsequently diluted to 1 M with 100 mM ammonium bicarbonate, trypsin was added at a 1:50 w/w ratio, and proteins were allowed to digest overnight at 37°C. The next day, neat LC-MS grade formic acid (Fisher Scientific) was added to a final concentration of 5% (v/v), prior to peptide desalting using C18 Spin Tips (100 µL, Pierce). Desalted peptides were dried in a vacuum concentrator, resolubilised in 50 μL of buffer A, and loaded onto a C18 trap column (Acclaim PepMap 100, 75 μm × 2 cm, 3 μm; Thermo Fisher Scientific). Trapped peptides were washed with 60 µL of Buffer A prior to loading onto a 15 cm Acclaim PepMap RSLC analytical column (75 μm × 15 cm, 2 μm, 100 Å, ThermoFisher) using a 45 min linear gradient of 5–40% buffer B (0.1% formic acid, 80% acetonitrile, 20% H_2_O, Fisher Scientific) at a flow rate of 300 nL min^-1^. Separated peptides were electrosprayed into an Orbitrap Eclipse Tribrid mass spectrometer in positive ion mode. The data-dependent MS^2^ spectra were collected as described above for lysates, but using a 60 min instrument method.

### In-gel digestion

Eluates from IPs were resolved on a 4-12% Bis-Tris SDS-PAGE gel (Pierce, Thermo Fisher Scientific) and stained with Coomassie Brilliant Blue. Gel bands of interest were excised using a fresh scalpel, diced into small ∼1 mm^3^ squares, and placed into a sterilised 1.5 mL Eppendorf tube. Gel pieces were destained using 50:50 100 mM ammonium bicarbonate:acetonitrile until no traces of Coomassie dye could be observed, and dried in a vacuum concentrator. Dried gel pieces were immersed in 20 mM TCEP (in 100 mM ammonium bicarbonate) and disulfide bonds were reduced by heating to 56°C for 1 hr. Residual TCEP solution was subsequently removed. Reduced cysteines were alkylated by soaking the gel pieces in 20 mM iodoacetamide (in 100 mM ammonium bicarbonate) in the dark for 1 hr. The gel pieces were washed twice with 50:50 100 mM ammonium bicarbonate:acetonitrile and dehydrated in a vacuum concentrator. The gel pieces were rehydrated with 20 ng uL^-1^ trypsin in 100 mM ammonium bicarbonate for overnight digestion at 37°C. The next day, the peptides trapped in the gel matrix were extracted twice by adding 200 µL of 47.5:47.5:5 acetonitrile:H_2_O:formic acid, and then once using 90:5:5 acetonitrile:H_2_O:formic acid. The solutions were combined in a fresh, sterilised tube and subsequently evaporated using a vacuum concentrator. Tryptic peptides were resuspended in 40 µL buffer A for LC-MS proteomic analysis. A 60 min gradient was used for separation of tryptic peptides on a 15 cm Acclaim PepMap RSLC analytical column. The peptides were electrosprayed into an Orbitrap Eclipse Tribrid mass spectrometer and data-dependent MS^2^ spectra were collected as described for above for immunoprecipitates.

### Electron transfer dissociation for glycopeptide identification

Gel bands for murine VGluT1 and mGluR2 were digested as described above for in-gel digestion. The tryptic peptides were loaded onto a C18 reverse phase trap column, washed with 40 μL of Buffer A at a flow rate of 20 μL min^-1^, and then loaded onto a 15 cm Acclaim PepMap RSLC analytical column (75 μm × 15 cm, 2 μm, 100 Å, Thermo Fisher Scientific) using a 40 min linear gradient of 5–40% buffer B (0.1% formic acid, 80% acetonitrile, 20% H_2_O, Fisher Scientific) at a flow rate of 300 nL min^-1^. After 40 min, the buffer composition was raised to 99% buffer B for 5 min prior to re-equilbration of the column at 2% buffer B for 4 min. Separated peptides were electrosprayed into an Orbitrap Eclipse Tribrid mass spectrometer in positive ion mode. MS^1^ spectra were collected at a resolving power of 120000 (at *m*/*z* 200). Peptide ions between charge state 2-8 above an intensity threshold of 5.0 × 10^4^ were selected using the quadrupole using an isolation window of *m*/*z* 2 and subjected to fragmentation using HCD (stepped normalised collision energy of 28, 30, and 32%). MS^2^ spectra were collected in the Orbitrap at a resolving power of 30000 (at *m*/*z* 200) and precursors within ±10 ppm were dynamically excluded for 60s. To trigger an electron transfer dissociation (with supplemental HCD) (EthcD) scan, precursor ions having HCD MS^2^ spectra containing diagnostic fragment ions at *m*/*z* 204.0867 (HexNAc), 138.0545 (HexNAc fragment), and 366.1396 (HexNAcHex) were reisolated using an isolation window of *m*/*z* 3, and subjected to EthcD (calibrated reaction parameters, supplemental HCD of 15%).

### Crosslinking MS (XL-MS) sample preparation, acquisition, and data analysis

Eluates were incubated with 3 mM DSSO (Pierce) for 30 min at room temperature before terminating the crosslinking reaction with 100 mM Tris, 0.02% DDM/0.002% CHS. The reaction was incubated for an additional ten minutes prior to unfolding in 4.5 M urea, reduction (with 10 mM TCEP), and alkylation (with 10 mM iodoacetamine), as described above for LC-MS based proteomic analysis of immunoprecipitates. Peptides were desalted using C18 peptide cleanup tips (Pierce).

Samples were analyzed using a NanoElute 2 UHPLC coupled to a timsTOF Ultra 2 mass spectrometer using a CaptiveSpray ionization source (Bruker). Peptides were separated on a 75 μm × 25 cm IonOptiks Aurora Ultimate C18 analytical column (IonOptiks). Buffer A was 0.1% FA in H_2_O and buffer B was 0.1% FA in LC-MS grade acetonitrile (ACN). A 30 or 120-minute linear gradient (0% to 40% buffer B) was used for elution.

Cross-linking data were acquired utilizing a DDA PASEF method with the following settings. Data were acquired in positive ionization mode over a scan range of 100 to 2500 m/z. TIMS settings were configured with an ion mobility range from 1/K0 = 0.60 Vs/cm^-2^ to 1.60 Vs/cm^-2^, and the ramp time was set to 100 ms to ensure optimal ion mobility separation. Collision energies were dependent on ion mobility values. To enhance modified peptide fragmentation, SE-CID settings were applied, with CE1 ranging from 1/K0 = 0.5 at 35 eV to 1/K0 = 1.6 at 54 eV, and CE2 ranging from 1/K0 = 0.5 at 40 eV to 1/K0 = 1.6 at 100 eV. The mass spectrometer was operated in PASEF mode, acquiring 6 PASEF MS/MS scans per cycle, with a target intensity of 20,000 and an intensity threshold of 500 to trigger MS/MS fragmentation. Active exclusion was enabled with precursors reconsidered after 0.40 minutes to avoid redundant fragmentation. Precursor ions with charge states between 3 to 8 were included for selection. A polygon filter was used in the ion mobility dimension to focus on target precursor ions. The MS/MS stepping list included collision RF values of 1600 Vpp for Scan #1 and 1200 Vpp for Scan #2, with corresponding transfer times of 100 µs and 64 µs, respectively. Additionally, pre-pulse storage times were set at 12 µs for Scan #1 and 10 µs for Scan #2.

Cross-linking data were searched using pLink (3.0.17), using an in-house curated protein database. The MS-cleavable flow was selected with DSSO as the linker used. Trypsin was selected with up to 3 missed cleavages allowed. Peptides with 3 or greater charge were analyzed. Carbimidomethylation was set as a fixed modification and methionine oxidation as dynamic. Identified cross-links were inspected in xiview.org.

### Glycoproteomic data analysis

Tryptic digests from gel bands corresponding to murine VGluT1 and mGluR2 were searched for both N- and O-linked glycopeptides using Byonic using the parameters described previously^93^. MS^2^ spectra were validated by manual inspection.

### LC-MS proteomic data analysis

Proteomic data were searched against the human or mouse proteomes (downloaded on 12/05/2023 and 09/03/2023, respectively, from Uniprot) in Maxquant (version 2.3.1.0) using default search parameters. Label-free quantification (LFQ) and intensity-based absolute quantification (iBAQ) were enabled. Custom FASTA files containing the amino acid sequences for Nb^VGluT1^ and Nb^mGluR2^ were also included during the searches of proteomic data of eluates from immunoprecipitation. N-terminal methionine loss and N-terminal acetylation were considered as variable modifications. Cysteine carbamidomethylation was set as a static modification. Perseus (version 2.0.11.0) was used for imputation^94^. Contaminants, reverse database hits, and proteins detected with only modified peptides, were removed. Values were log2 transformed and missing protein abundances observed in >70% of the biological replicates were imputed using the default settings. All data matrices were imported to OriginPro 2021 for plotting.

### Native MS-guided lipidomic analysis

To explore the lipid microenvironment of murine mGluR2, 12 µL of eluate from n=7 biologically independent immunoprecipitations were processed via our native MS-guided lipidomic pipeline^56^. Briefly, eluates were supplemented with 1 µL SPLASH lipidomix (Avanti Polar Lipids), and lipids were extracted using the methyl-*tert*-butyl ether (MTBE):methanol approach^95^. The mixture was mixed with 300 µL methanol and 1 mL of MTBE, and incubated at room temperature for one hour. 250 µL of H_2_O was then added, the mixture was vortexed vigorously, then allowed to sit undisturbed to promote phase separation. The mixture was centrifuged at 400×*g* for 10 min at room temperature, and the organic phase was transferred to a new vial. The extraction was repeated with 400 µL of the extraction mixture (10/3/2.5 of MTBE/methanol/H_2_O). The resulting organic phases were combined and dried under nitrogen in a fume hood. The lipid films were resuspended in 40 µL of 80% mobile phase A (acetonitrile/H_2_O [60/40, vol/vol] supplemented with 10 mM ammonium formate and 0.1% [vol/vol] formic acid) and 20% mobile phase B (isopropanol/acetonitrile [90/10, vol/vol] supplemented with 10 mM ammonium formate and 0.1% [vol/vol] formic acid). The resuspended lipids were sonicated for 5 min in a bath sonicator and transferred to a glass vial for analysis. For whole brain or OFC lysate bulk lipidomics, lipids were extracted as described above from 100 µg of total protein.

1 µL of lipids were separated on a 15 cm Acclaim PepMap RSLC C18 reverse phase analytical column (75 μm × 15 cm, 2 μm, 100 Å, ThermoFisher) using an Ultimate 3000. Lipids were electrosprayed into an Orbitrap Eclipse in both positive and negative modes. Instrument parameters and chromatography gradients are described in Ref ^56^.

### Native mass spectrometry of endogenous and recombinant mGluRs

An Orbitrap UHMR mass spectrometer, tuned to transmit high *m*/*z* ratio ions ^96^, was used to acquire native mass spectra for mGluRs (human, mouse, and recombinant). Proteins were buffer exchanged into 400 mM ammonium acetate (pH 7.4) supplemented with 0.17%/0.017% n-decyl-β-D-maltoside (DM)/CHS twice using Pierce Zeba desalting columns (75 µL, 40 kDa MWCO). 1–3 µL of buffer exchanged sample was loaded into gold-coated nanoelectrospray capillaries prepared in-house. All spectra were collected in the positive ion mode. Typical instrument parameters included: +800–1200 V electrospray bias; capillary temperature 200–300°C; 200–240 V desolvation voltage (in-source trapping); 50–75 V source collision induced dissociation voltage; 220 – 270 V HCD voltage. Typically, a resolution of 25000 (at *m/z* 200) was used. Prior to MS analysis of immunoprecipitated mGluR2, instrument parameters were first optimised using recombinant mGluR2. We observed limited drift in required parameters (±20 V HCD and source parameters) over the course of the study.

### Native mass spectrometry of VGluT1

A modified Orbitrap Ascend Tribrid mass spectrometer^31^ was used to generate native mass spectra of VGluT1 (mouse, human, recombinant). VGluT1 was buffer exchanged twice into 400 mM ammonium acetate (pH 7.4) supplemented with 0.17%/0.017% DM/CHS using Pierce Zeba desalting columns (75 µL, 40 kDa MWCO). The instrument was operated in “high-pressure, intact protein” mode and data were collected in the positive ion mode. 1–3 µL of VGluT1 was introduced into the mass spectrometer using gold-coated nanoelectrospray capillaries prepared in-house. Typical instrument parameters: 800–1100 V electrospray bias; 175–200 V in-source dissociation bias; source CID compensation scaling factor 0.01–0.04; 100 ms injection time and 100% AGC target with a maximum injection time of 500 ms. Data were collected in the Orbitrap mass analyzer at a resolving power of 15000 at *m/z* 200. VGluT1 was further liberated from DM/CHS detergent micelles using 10 ms of infrared irradiation at 6–8% output power (of a 60W Ti Firestar 10.6 µm laser) in the high-pressure region of the linear ion trap.

### Native mass spectrometry data analysis

Native mass spectra were extracted from QualBrowser (Xcalibur version 4.1.31.9) as text files and imported into OriginPro 2021 (v9.8.0.200). Spectra were processed via adjacent averaging smoothing with a 10-point window size and normalized between 0 and 1 for plotting.

### Proton Transfer Charge Reduction

The charge state peaks in native mass spectra of mGluR2 were broad, making it difficult to assign the charge state (and therefore, mass) based on peak spacing alone. To determine charge states, therefore we used the ion trap to isolate a single peak in the native mass spectrum of mGluR2 (25–50 *m*/*z* wide) and carried out a proton transfer reaction to generate a distribution of charge reduced product ions^97^. Adjacent charge states of these product ions can be readily resolved, allowing for algebraic determination of their charge states based on spacing. Typical ion-ion reaction parameters were reagent target of 1 × 10^6^ and 5–6 ms reaction time.

### Top-down mass spectrometry data collection and analysis

Top-down native mass spectrometry experiments were conducted on a modified Orbitrap Ascend Tribrid mass spectrometer which included the addition of a 10.6 µm wavelength CO_2_ laser (Synrad Firestar Ti60 CO_2_ continuous wave IR laser) directed into the high-pressure cell of the linear ion trap analyser. The timing and power of the laser output were controlled using modified instrument software^31^. The instrument was operated in high-pressure mode (20 mTorr in front and back IRMs). Typical source parameters were: 900-1100 V electrospray bias, 150°C transfer tube temperature, RF lens 150%. The source CID voltage and compensation scaling factors were optimized for each protein: mGluR2 utilized 250 V source CID and a compensation scaling factor of 0.05, and VGluT1 required 180 V source CID and a compensation scaling factor of 0.03. Source CID compensation is a feature available on Tribrid mass spectrometers, which serves to improve transmission of proteins and complexes through kinetic energy management, and has been described elsewhere ^98^. To generate MS^1^ spectra, the ions were subjected to a 5 ms burst of infrared irradiation (at 7 – 9% of the maximum output power, or ca. 4.2 – 5.4 W) before detection in the Orbitrap at a resolution of 15000 @ *m*/*z* 200.

For MS^2^ experiments, ions within an *m*/*z* region of interest were isolated in the ion trap using a selection window 50–100 *m*/*z* wide (VGluT1) or 100–300 *m*/*z* wide (mGluR2) and a q-value of 0.1. Isolated ion populations were subjected to infrared multiphoton dissociation (IRMPD) for 10 ms using 12% of the maximum laser output (ca. 7.2 W) to induce fragmentation. Fragment ions were detected in the Orbitrap mass analyzer using a resolving power of 240,000 at *m/z* 200. Maximum injection times were set to 1 s (with a 100% AGC target) and averaged for maximum number of individual scans (999 scans). For non-covalent lipid release, peaks were isolated in the linear ion trap and the lipids released by <100 V HCD. Lipids were detected in the ion trap (“Turbo” scan mode).

Fragments were manually validated against the UniProt sequences of mGluR2 (UniProt ID: Q14416), mGluR3 (UniProt ID: Q14832), and VGluT1 (UniProt ID: Q9P2U7) using precisION.^38^ The characteristic fragments for mGluR2 and mGluR3 were further confirmed by MS^3^ analyses. Briefly, fragments at *m*/*z* 1693 and *m*/*z* 1751, were individually isolated in the ion trap using a 5 – 10 *m*/*z* isolation window. Fragment ions were generated using higher-energy collisional dissociation (HCD) (20 – 40 normalized energy) followed by detection in the ion trap using the “Turbo” scan mode. MS^3^ data were manually annotated. Plots showing fragmentation depth and additional analyses were carried out using precisION.^38^ We could not generate nTDMS spectra (S/N > 3) for mGluR2 immunoprecipitates from CON8 and DEPR7.

### Scaling factors of top-down data of human and mouse mGluR2/3

Recombinant murine and human mGluR2/3 complexes were purified from FACS-enriched stably-transfected HEK293 GNTI^-/-^ cells via tandem affinity purification (FLAG on mGluR2, StrepII on mGluR3). mGluR2/3 contain a 1:1 stoichiometry between mGluR2 and mGluR3. In nTDMS studies of these complexes, we noted differences in the ion yields for mGluR2- and mGluR3-derived fragments; mGluR2 fragment ion series’ consistently dominated the nTDMS spectra. We therefore established an empirically-determined correction factor to scale the intensities of mGluR3 fragment ions to adjust for the expectation that these fragments *must* emerge from mGluR2/3 (having subunits present at molar equivalent levels). We restricted the quantification to fragments that arose from structurally-equivalent sites in the extracellular domains of the mGluR2 and mGluR3 protomers, and were consistently observed in endogenous complexes. After application of the calibrated scaling factor, all of the relative mGluR3-fragment intensity reports on mGluR2/3 heterodimers. The residual mGluR2 intensity reports on mGluR2/2 homodimers.

### XL-MS guided HADDOCK docking

Atomic coordinates for the mGluR2/3 heterodimer were obtained from the cryo-EM structure of mGluR2/3 heterodimer in the active conformation (8JD2).^57^ The extracellular domain (ECD) was extracted for each subunit. Atomic coordinates for CRMP2 were obtained from the crystal structure of human CRMP2 (PDB: 5X1A),^66^ with the biological assembly generated to produce the functional homotetramer. All structure preparation steps were in Python.

Solvent-accessible surface area (SASA) was calculated for each prepared structure using the Shrake-Rupley algorithm as implemented in Biopython, with per-residue relative SASA computed against maximum theoretical values as described.^99^ Residues with relative SASA ≥ 25% were classified as surface-exposed. The experimentally identified DSSO crosslink between mGluR3 K423 and CRMP2 K398 was used to define the docking interface. mGluR3 K423 and CRMP2 K398 were designated as active residues, with passive residues defined as surface-exposed residues within 12 Å of each active residue as determined by Cα-Cα distance calculations.

Data-driven docking was performed using the HADDOCK 2.4 web server^73^ with Guru-level access. The mGluR2/3 ECD heterodimer was submitted as molecule 1 and the CRMP2 tetramer as molecule 2. The DSSO crosslink was incorporated as an unambiguous distance restraint with an upper bound of 35 Å, corresponding to the maximum Cα-Cα distance achievable with the DSSO spacer arm of 10.1 Å accounting for side chain flexibility. Ambiguous interaction restraints (AIRs) were generated from the active and passive residue definitions following the standard HADDOCK AIR protocol.^72^ The “remove buried active/passive residues” filter was disabled to retain the experimentally validated crosslink restraint at CRMP2 K398. Default HADDOCK docking parameters were used, generating 1000 rigid body docking structures, of which 200 were selected for semi-flexible refinement and 200 for final explicit solvent refinement. Structures were clustered by RMSD with a 7.5 Å cutoff. The top-scoring cluster was selected based on HADDOCK score, Z-score, and restraint violation energy.

The best-scoring docked structure from cluster 1 (HADDOCK score −94.2 ± 8.5, Z-score −1.8) was selected for further analysis. The full-length mGluR2/3 structure including transmembrane domains was reconstructed by superimposing the docked ECD complex onto the corresponding chains of PDB 8JD2 using the matchmaker function in UCSF ChimeraX,^100^ with RMSD values of 0.284 Å and 0.279 Å for mGluR2 and mGluR3 respectively. The CRMP2 homotetramer was reconstructed by superimposing the biological assembly of PDB 5X1A onto the docked CRMP2 monomer using ChimeraX matchmaker (RMSD 0.347 Å), confirming correct placement of the tetramer relative to the mGluR2/3 ECD. The Cα-Cα distance between the crosslinked residues mGluR3 K423 and CRMP2 K398 in the final model was 11.9 Å, consistent with the experimentally observed DSSO crosslink.

### Rendering CryoEM structures

All structures were rendered in ChimeraX (UCSF)^100^.

