## SupplementaryInformation for "Molecular dissection of protein complexes isolated from sections of human brain"

### List of supplemental figures

**Figure S1.** Proton transfer charge reduction to determine the charge states and therefore mass of murine mGluR2
**Figure S2** Native mass spectra of recombinant mGluR2/2 homodimers bound to $Nb^{mGluR2}$
**Figure S3** Native mass spectra of recombinant mGluR3/3 homodimers with to $Nb^{mGluR2}$
**Figure S4** Native mass spectra of recombinant mGluR2/3 heterodimers bound to $Nb^{mGluR2}$
**Figure S5.** Native top down mass spectrum of palmitoylated mGluR3 fragment ions
**Figure S6**. Proteomic analysis of iPSC-derived glial cell conditioned
**Figure S7.** $MS^3$ determination of the amino acid composition of recombinant mGluR2 and recombinant mGluR3 fragments
**Figure S8**. The impact of post-mortem interval (PMI) on the integrity of endogenous human VGluT1 and mGluR2.
**Figure S9.** Comparison of representative native top-down MS spectra of intact human VGluT1 from CON and DEPR groups
**Table S1.** Cohort details from sgACC
**Table S2.** Cohort details from OFC
**Table S3**. Proteomic analysis of the ~45 kDa band in Fig 4A.

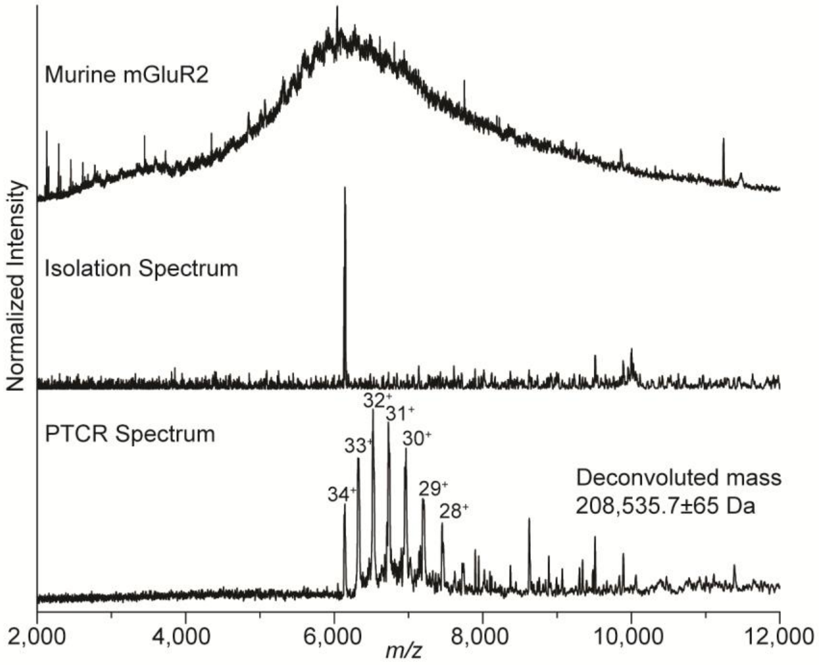


**Figure S1. Proton transfer charge reduction to determine the charge states and therefore mass of murine mGluR2.** (Top) a native mass spectrum of mGluR2 from a C57BL6 murine brain. Algebraic determination of charge states for each m/z peak results in charge ambiguity (±1 *z* leading to uncertainty in mass). (Middle) a spectrum resulting from the isolation of a single charge state peak (ion trap isolation, ±12.5 *m*/*z* window). (bottom) a spectrum recorded after the generation of charge reduced product ion peaks via proton transfer-charge reduction (PTCR). Isolation and charge reduction of a single peak (from the top spectrum) enables generation of well-spaced peaks with defined widths (bottom spectrum), making it possible to determine algebraically the charge states based on peak spacing. Therefore, determination of the mass is readily possible.

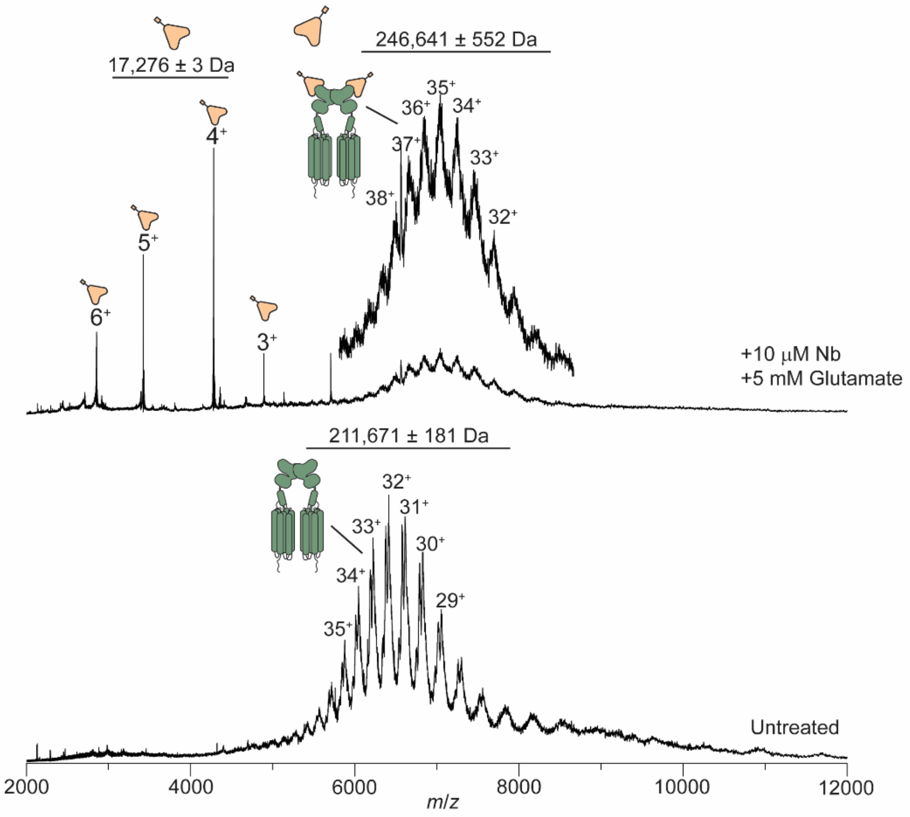


**Figure S2 Native mass spectra of recombinant mGluR2/2 homodimers and nanobody binding following incubation with an excess of Nb$^{mGluR2}$ and excess glutamate.** The measured masses are consistent with recombinant mGluR2/2 homodimers and two copies of Nb$^{mGluR2}$ binding to each mGluR2/2 protomer.

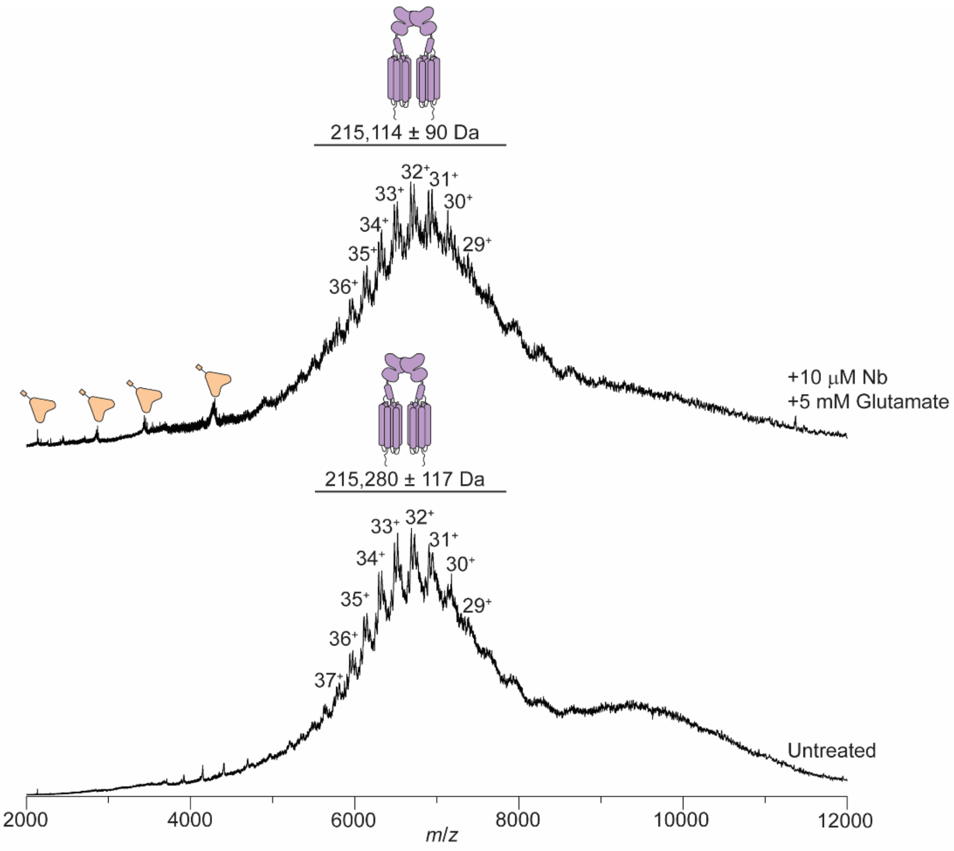


**Figure S3. Native mass spectra of recombinant mGluR3/3 homodimers and nanobody binding following incubation with an excess of Nb$^{mGluR2}$ and excess glutamate.** The measured mass is consistent with recombinant mGluR3/3 homodimers and no binding of Nb$^{mGluR2}$ to mGluR3/3 was observed.

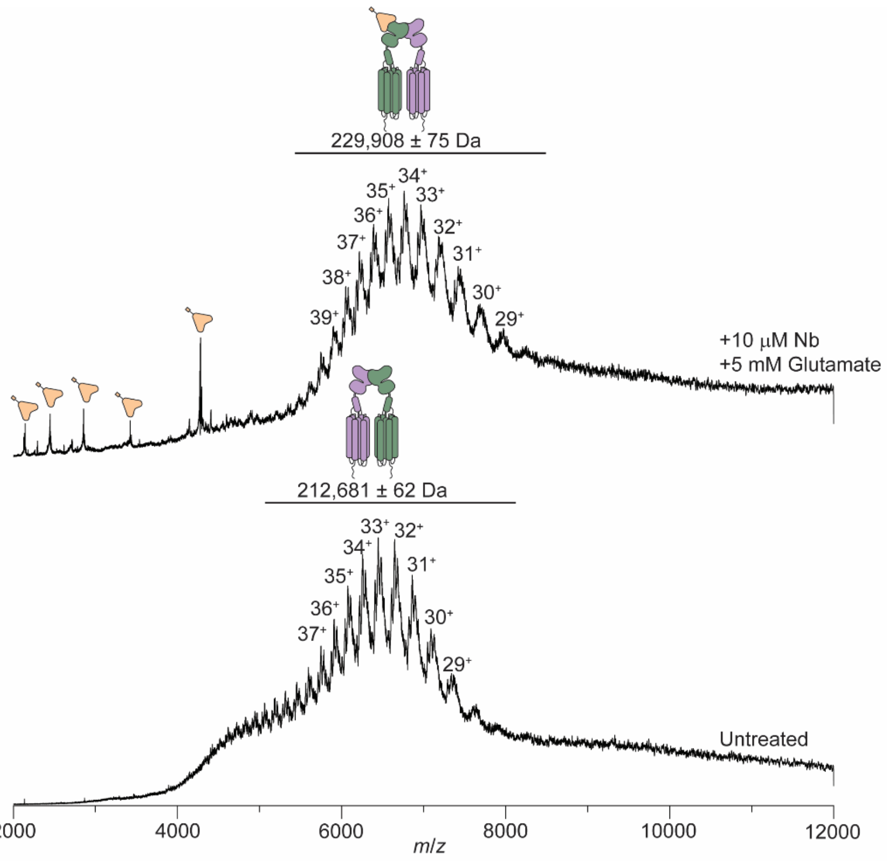


**Figure S4. Native mass spectra of recombinant mGluR2/3 heterodimers and with nanobody binding.** (Bottom) Native mass spectrum of recombinant mGluR2/3 purified. The mass is consistent with mGluR2/3 binding in a 1:1 stoichiometry. (Top) Native mass spectrum of recombinant mGluR2/3 following incubation with excess of $Nb^{mGluR2}$ and excess glutamate. The measured mass is consistent with one copy of $Nb^{mGluR2}$ binding to recombinant mGluR2/3 heterodimers.

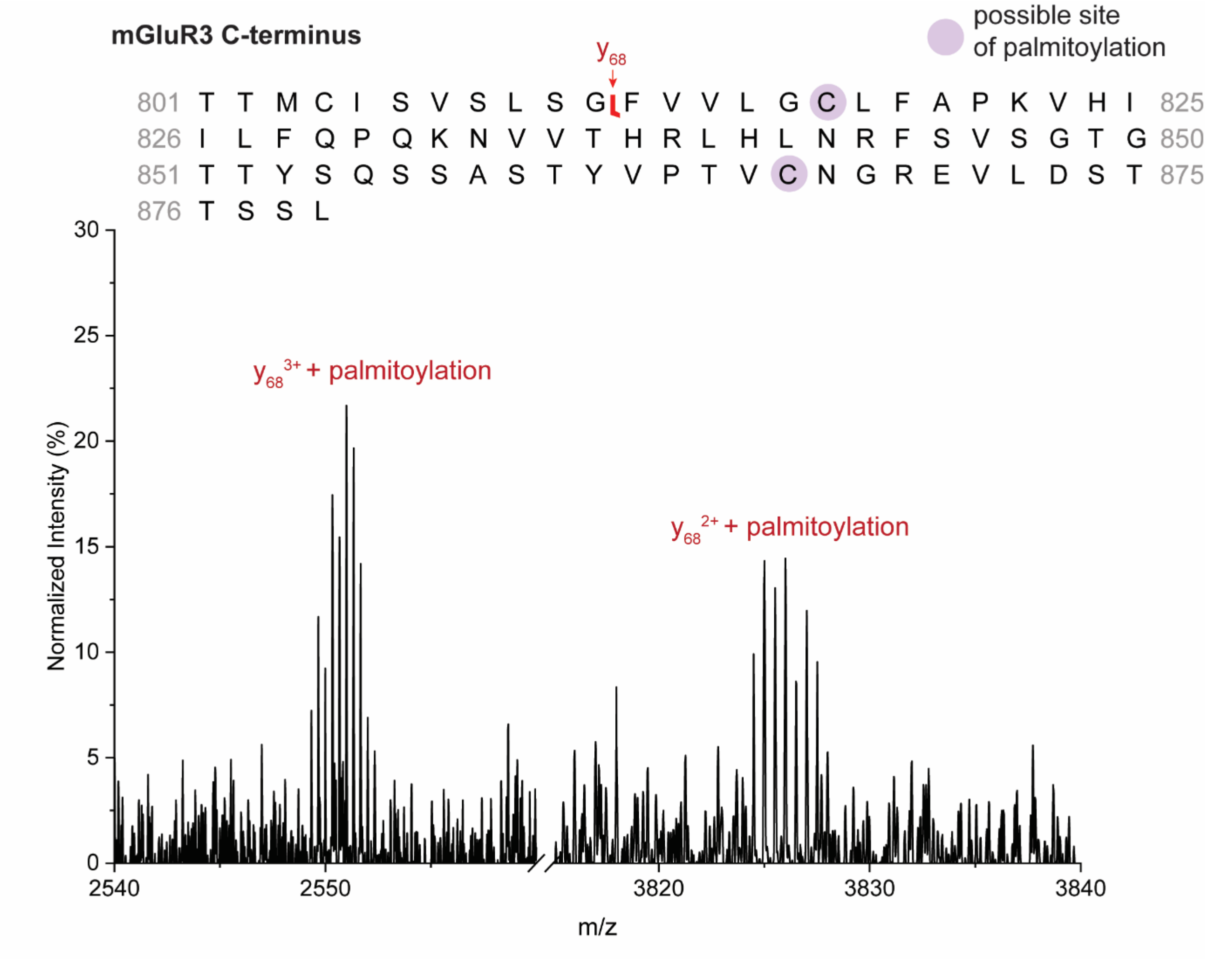


**Figure S5. nTDMS spectrum of palmitoylated fragments on mGluR3 in endogenous mGluR2/3 from OFC.** Only one palmitoylation modification was attached to y68 fragment derived from the C-terminus of mGluR3. This PTM could be present at either of the two cysteine residues indicated in purple. Fragment ion sequences were determined using precisION.

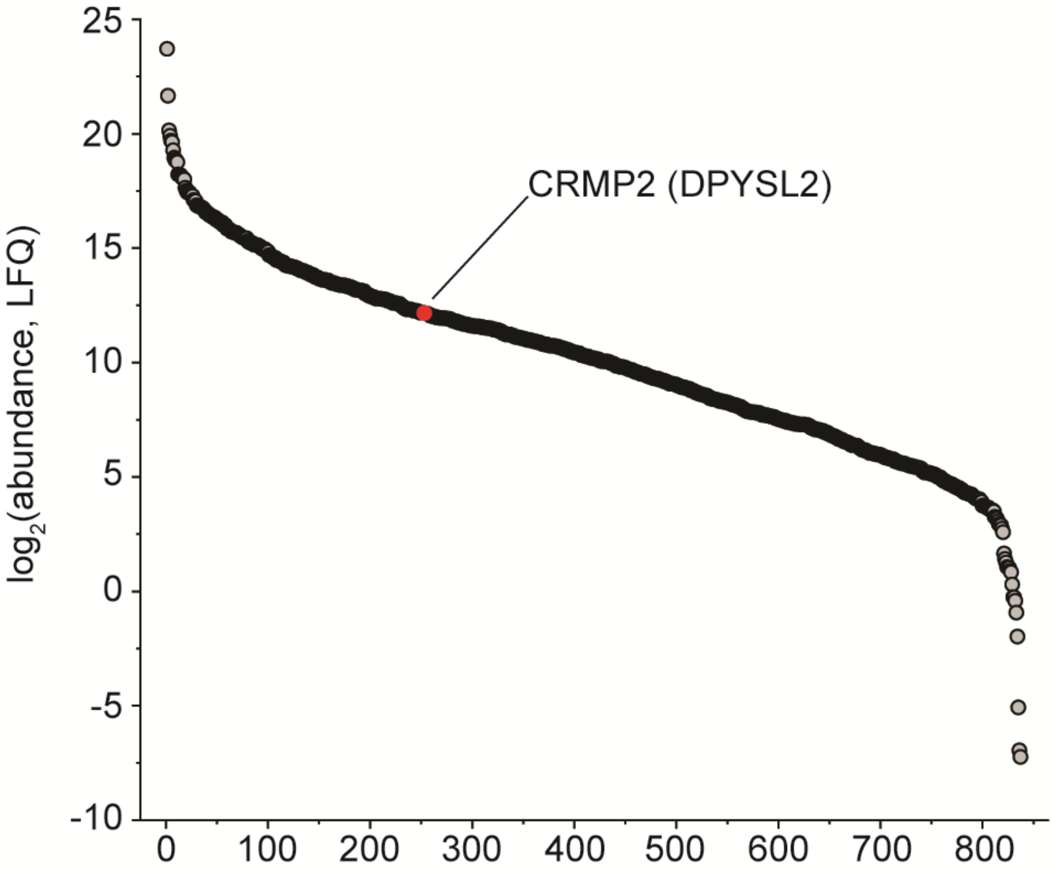


**Figure S6. Proteomic analysis of iPSC-derived glial cell conditioned media (from Trindade et al Mol. Psychiatry. 2022).** Plot of $\log_2$(abundance) relative to protein rank for the proteins that are secreted from iPSC-derived glia. Red circle indicates CRMP2 (gene name DPYSL2). Data replotted from Trindade, P., Nascimento, J.M., Casas, B.S. et al. Induced pluripotent stem cell-derived astrocytes from patients with schizophrenia exhibit an inflammatory phenotype that affects vascularization. Mol Psychiatry 28, 871–882 (2023). https://doi.org/10.1038/s41380-022-01830-1.

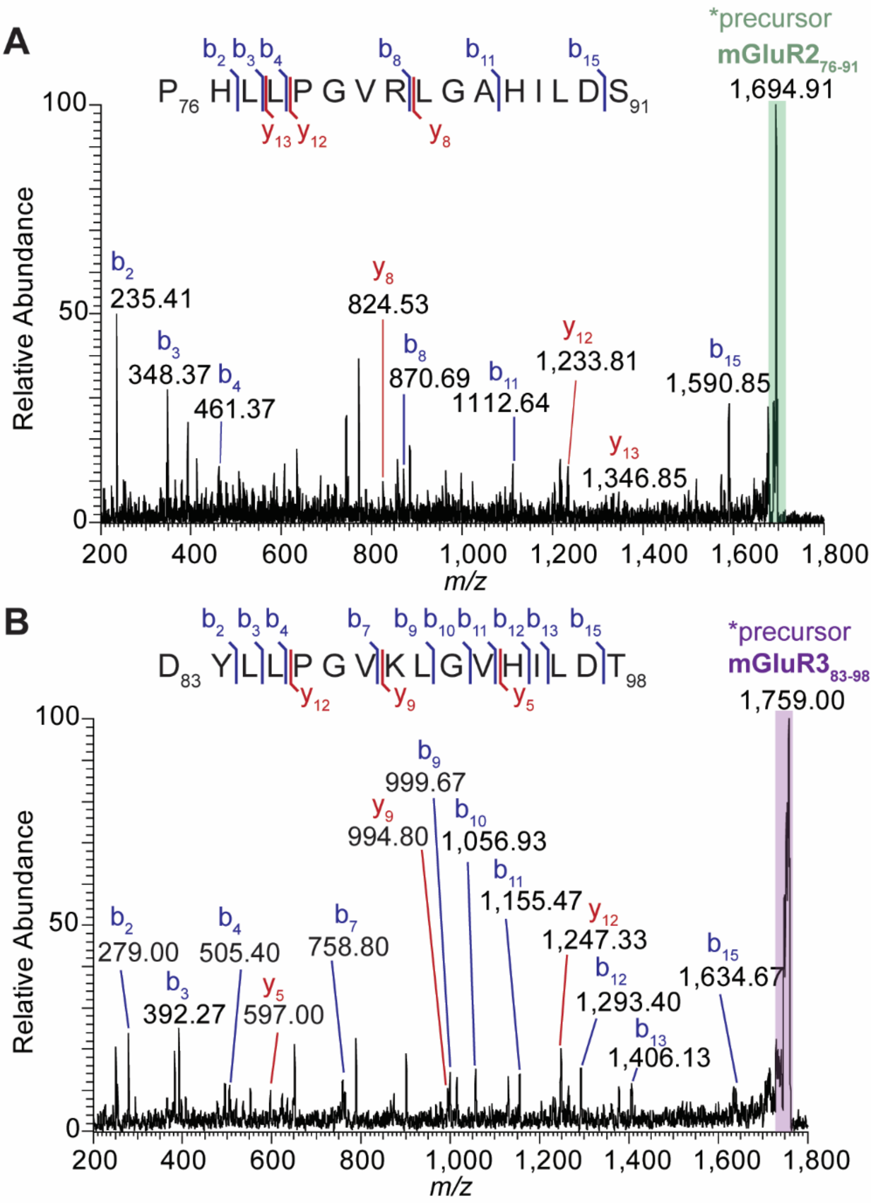


**Figure S7**. **MS$^3$ determination of the amino acid composition of recombinant mGluR2 and recombinant mGluR3 fragments at *m*/*z* 1693 and *m*/*z* 1751, respectively.** (A) MS$^3$ isolation and fragmentation of the mGluR2 fragment at *m*/*z* 1693 (isolation width of 10 *m*/*z*) revealed b-type and y-type ions consistent with the 16 amino acids between residues 76-91 of mGluR2. (B) MS$^3$ isolation and fragmentation of the mGluR3 fragment at *m*/*z* 1751 (isolation width of 10 *m*/*z*) revealed b-type and y-type ions consistent with the corresponding 16 amino acids between residues 83-98 of mGluR3.

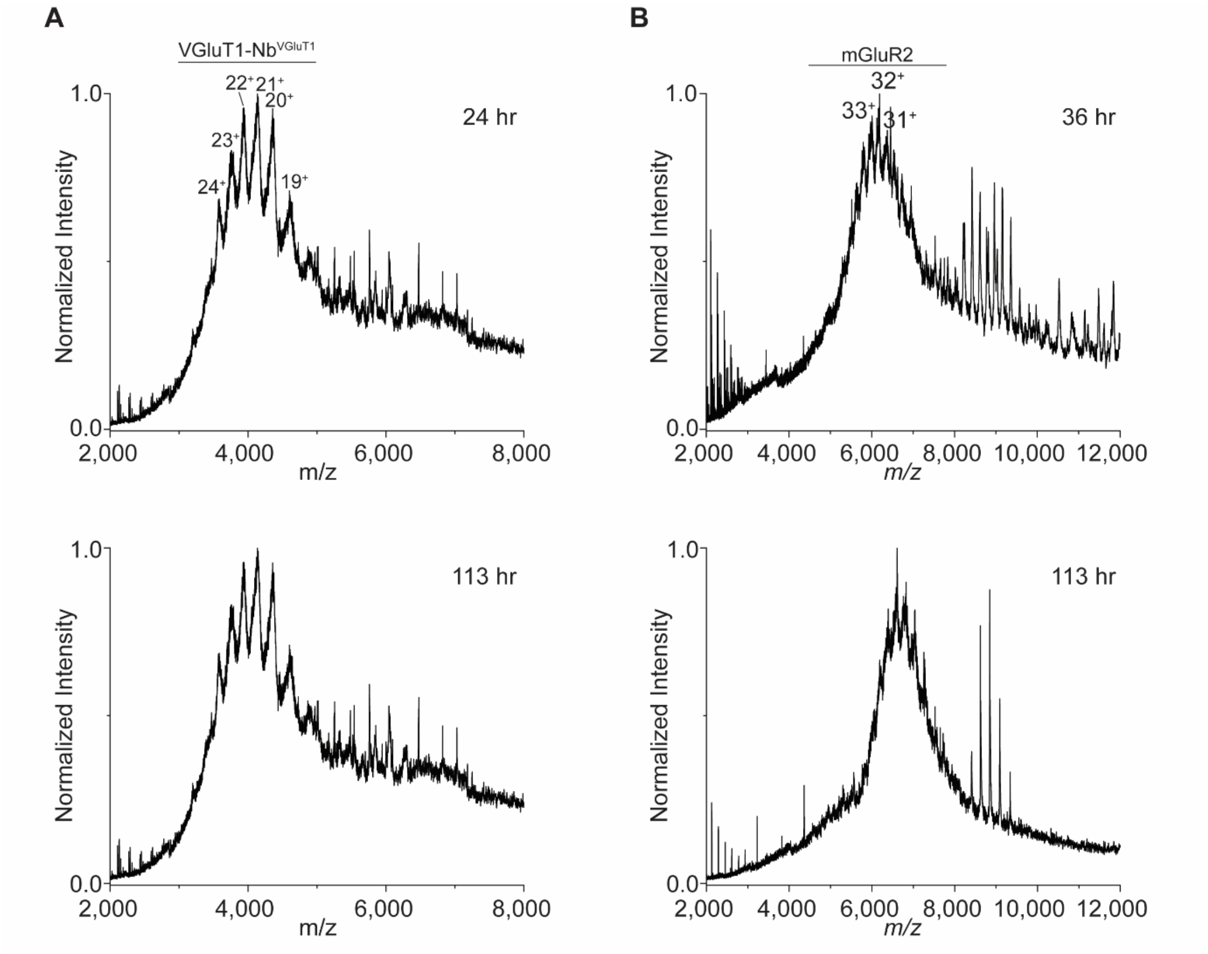


**Figure S8. The impact of post-mortem interval (PMI) on the integrity of endogenous human VGluT1 and mGluR2.** (A) Native mass spectra of VGluT1 immunoprecipitated from two healthy control subjects at different post-mortem intervals. (B) native mass spectra of mGluR2 from two healthy control subjects at different PMIs. Tissue was harvested at well-defined PMIs from human OFC by a neuropathologist. For both VGluT1 and mGluR2, no apparent differences in spectral characteristics for each respective protein complex could be correlated with PMI, e.g., no clear trends were observed in measured m/z (mass), presence of high charge states due to unfolding, new interactors, or emergence of mGluR2 monomers.

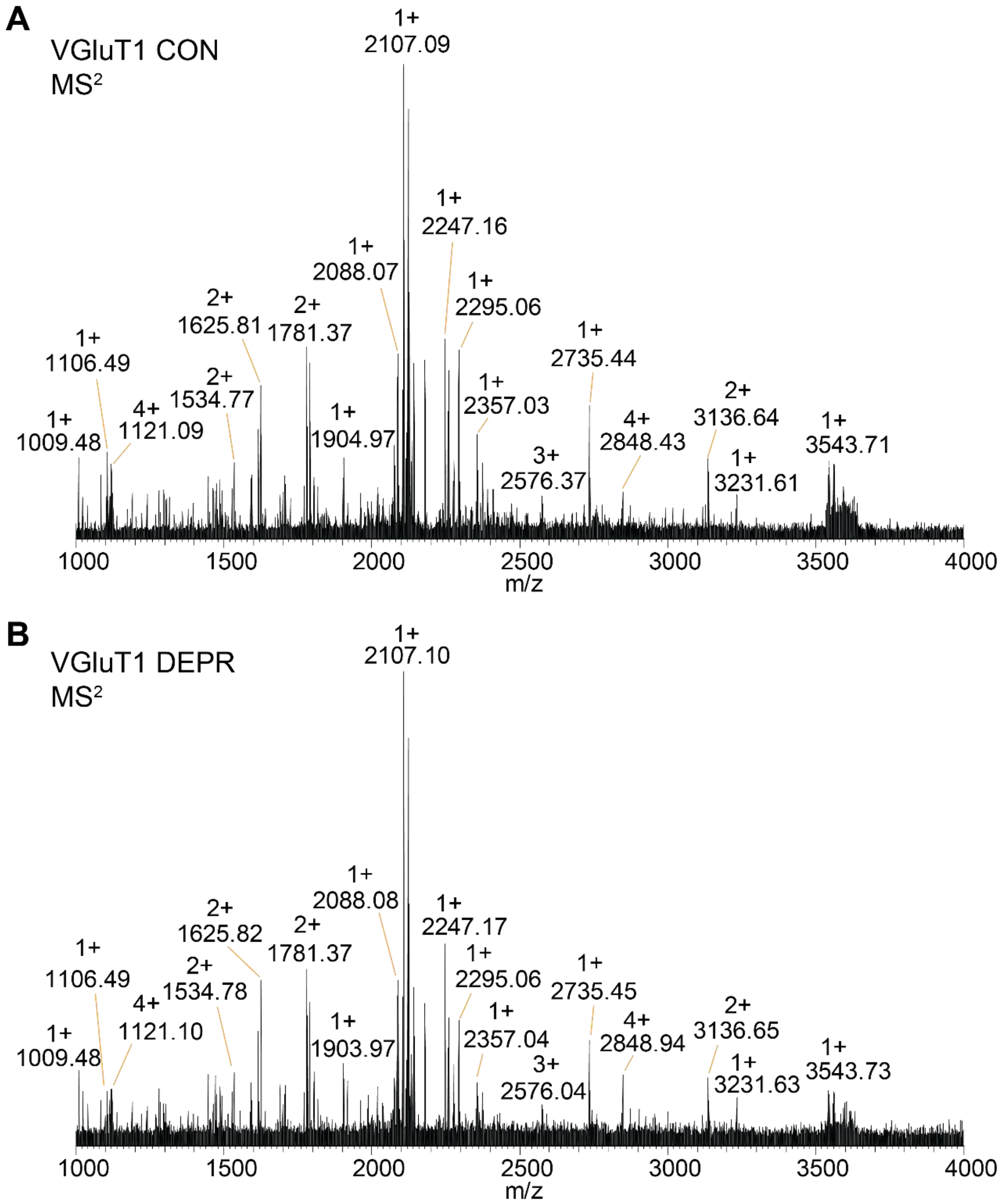


**Figure S9. Comparison of representative native top-down MS spectra of intact human VGluT1 from CON and DEPR groups.** Isolation of *m/z* peaks assigned to the intact complex of VGluT1 from (A) CON and (B) DEPR shows no apparent differences, suggesting the integrity of the transporter is not distinguishable at this level.

**Table S1. Details of sgACC donors used in the study**

| Case number on UK Brain bank network database | Internal Oxford Brain Bank case ID | Brains for Dementia Research case number | Age | Sex | Neuropath diagnosis | Year of death | Geriatric Depression Scale (GDS score closest to death) |
|---|---|---|---|---|---|---|---|
| BBN004.31477 | 17/109 | O-0473 | 77 | F | Normal aged brain (Braak I, Thal phase 0) | 2017 | 0 (2016) |
| BBN004.37449 | 22/061 | O-T180/0238, N0238 | 79 | M | Normal brain with some age-related pathology (Braak I, Thal phase 3). | 2022 | 7 (2022) |
| BBN004.37446 | 22/033 | O-0110 | 85 | F | Normal brain with some age-related AD pathology (Braak II, Thal phase 1) | 2022 | 1 (2021) |
| BBN004.37443 | 22/024 | O-T132/0172, N0172 | 82 | M | Normal brain with some age-related AD pathology (Braak I, Thal phase 0) | 2022 | 3 (2019) |

**Table S2. Details of OFC donor subjects used in the study**

| Subject Number | Autopsy ID | Age | Gender | PMI (hr)[a] | Cause of Death | Past Medical History | Past Prescription History |
|---|---|---|---|---|---|---|---|
| CON 1 | SD024/14 | 38 | M | 36 | 1a) Ischaemic heart disease<br>1b) Coronary artery atherosclerosis<br>2) Type 2 diabetes | Type 2 diabetes<br>Hypertension<br>Depression<br>Hypercholesterolaemia<br>Smoker | Lisinopril 20 mg<br>Metformin 1G<br>Gliclazide 160 mg<br>Pravastatin 40 mg<br>Amitriptyline 150 mg<br>Dihydrocodeine |
| CON 2 | SD026/16 | 37 | F | 126 | 1a) Myocardial fibrosis (cause uncertain) | Crohn's disease<br>Non-smoker<br>Minimal alcohol | Nil |
| CON 3 | SD008/14 | 43 | M | 96 | 1a) Coronary artery thrombosis<br>1b) Coronary artery atherosclerosis | Smoker | Nil |
| CON 4 | SD010/15 | 57 | F | 113 | 1a) Coronary artery atheroma | Non-smoker | Nil |

| | | | | | | | |
|---|---|---|---|---|---|---|---|
| CON 5 | SD014/18 | 46 | F | 99 | 1a) Complications of coronary artery dissection and myocardial scarring | Depression<br>Smoker<br>Minimal alcohol | Venlafaxine 75 mg<br>Hydroxyzine 25 mg |
| CON 6 | SD038/17 | 34 | M | 99 | 1a) Ischaemic heart disease<br>1b) Coronary artery atheroclerosis | Non-smoker<br>Minimal alcohol | Nil |
| CON 7 | SD042/18 | 73 | F | 74 | 1a) Hypertensive heart disease | Hypertension<br>Osteoarthritis<br>Previous smoker | Unknown |
| CON 8 | SD005/15 | 46 | F | 76 | 1a) Complications of ischaemic heart disease and hepatic steatosis<br>2) Obesity | Depression<br>OCD<br>Hypothyroidism<br>Anxiety<br>Smoker | Diazepam 5 mg<br>Thyroxine 25 mcg |
| DEPR 1 | SD016/13 | 44 | F | 70 | Suicide | Type 1 diabetes<br>Depression | Simvastatin 40 mg<br>Montelukast 10 mg<br>Venlafaxine 75 mg<br>Trazodone 150 mg<br>Diazepam 2 mg<br>Citalopram 40 mg<br>Fluoxetine 20 |

| | | | | | | | |
|---|---|---|---|---|---|---|---|
| | | | | | | | mg<br>Seretide inhaler |
| DEPR 2 | SD048/12 | 63 | M | 44 | Suicide | Depression<br>Melanoma | Diazepam 2 mg<br>Nortriptyline 140 mg<br>Risperidone 1 mg |
| DEPR 3 | SD036/09 | 44 | F | 49 | Suicide | Depression<br>Smoker<br>Minimal alcohol<br>Previous overdoses<br>Migraine | Propanolol 40 mg<br>Paroxetine 30 mg |
| DEPR 4 | SD039/08 | 41 | M | 41 | Suicide | Schizophrenia<br>Depression<br>Anxiety | Paroxetine 25 mg |
| DEPR 5 | SD046/16 | 42 | M | 103 | Suicide | Anxiety<br>Depression<br>Alcohol misuse<br>Polysubstance abuse<br>Smoker | Imipramine 25 mg |
| DEPR 6 | SD009/06 | 24 | F | 88 | Suicide | Depression | Fluoxetine 20 mg |

| | | | | | | | |
|---|---|---|---|---|---|---|---|
| DEPR 7 | SD010/10 | 39 | M | 81 | Suicide | Depression | Venlafaxine 150 mg |
| a. PMI is determined from the time subjects were found, not the time of death, and is therefore an estimate. | | | | | | | |

**Table S3. List of proteins found in the band corresponding to glutamine synthetase in Fig 4A**

| Protein names | Gene names | MW (kDa) | log2(iBAQ) |
|---|---|---|---|
| Glutamine synthetase | GLUL | 57.174 | 27.921 |
| Pyruvate dehydrogenase E1 | PDHA1 | 46.396 | 20.932 |
| Actin, cytoplasmic 2 (multiple forms) | ACTG1;ACTB;POTEF;ACTA2;ACTA1;ACTG2;ACTC1 | 38.975 | 20.537 |
| Keratinocyte proline-rich protein | KPRP | 64.135 | 20.224 |
| Myelin proteolipid protein | PLP1 | 30.077 | 18.626 |
